# Ultrastable and Versatile FoxP3 Ensembles on Microsatellites

**DOI:** 10.1101/2024.12.12.628245

**Authors:** Fangwei Leng, Raquel Merino, Xi Wang, Wenxiang Zhang, Taekjip Ha, Sun Hur

**Affiliations:** Howard Hughes Medical Institute and Program in Cellular and Molecular Medicine, Boston Children’s Hospital, MA 02115, USA; Department of Biological Chemistry and Molecular Pharmacology, Harvard Medical School, Boston, MA 02115, USA; Department of of Biology, Johns Hopkins University, Baltimore, MD 21218

## Abstract

Microsatellites are essential genomic components increasingly linked to transcriptional regulation. FoxP3, a transcription factor critical for regulatory T cell (Treg) development, recognizes TTTG repeat microsatellites by forming multimers along DNA. However, FoxP3 also binds a broader range of TnG repeats (n=2-5), often at the edges of accessible chromatin regions. This raises questions about how FoxP3 adapts to sequence variability and the potential role of nucleosomes. Using cryo-electron microscopy and single-molecule analyses, we show that FoxP3 assembles into distinct supramolecular structures depending on DNA sequence. This structural plasticity enables FoxP3 to bridge 2-4 DNA duplexes, forming ultrastable structures that coordinate multiple genomic loci. Nucleosomes further facilitate FoxP3 assembly by inducing local DNA bending, creating a nucleus that recruits distal DNA elements through multiway bridging. Our findings thus reveal FoxP3’s unusual ability to shapeshift to accommodate evolutionarily dynamic microsatellites and its potential to reinforce chromatin boundaries and three-dimensional genomic architecture.

## INTRODUCTION

Microsatellites, tandem repeats of short sequences (1-6 nt), constitute approximately 5% of the human genome^1,2^. Traditionally, they have been regarded as incidental genetic components with high mutability, contributing to interindividual genetic variability^3,4^ and, in some cases, to disease by disrupting existing genetic elements^1,2,5^. However, recent evidence suggests that microsatellites may play direct and positive roles in transcriptional regulation^6–11^. Many microsatellites are evolutionarily enriched in gene-proximal cis-regulatory elements^9,12,13^, and approximately 90% of transcription factors (TFs) tested, in particular those with the forkhead DNA-binding domain (DBD), have demonstrated the ability to bind specific microsatellites both in vitro and in vivo for their transcriptional functions^6,14,15^. Despite the growing interest in these interactions, the mechanisms by which TFs recognize microsatellites remain largely unclear. The heterogeneous and evolutionarily dynamic nature of microsatellites points to novel principles yet to be uncovered in TF-microsatellite interactions.

FoxP3 is a central TF in the development and function of regulatory T cells (Tregs), a subset of T cells crucial for maintaining immune homeostasis^16–18^. Loss-of-function mutations or knockout of FoxP3 results in severe autoimmunity in both humans and mice^17,19–22^. FoxP3 is a forkhead TF and was long thought to recognize the forkhead consensus motif TGTTTAC, namely FKHM^23,24^. Structural and biochemical studies later revealed that FoxP3 preferentially binds to inverted-repeat FKHM (IR-FKHM) sequences, forming a head-to-head dimer^25^. However, these sequences—whether single-FKHM or IR-FKHM—have not been strongly supported as the primary genomic targets of FoxP3 in Tregs^25^. Instead, recent studies have identified TnG repeat microsatellites (n=2-5) as the primary drivers of FoxP3’s genomic occupancy, showing high-affinity binding to these sequences^14,15^. The cryo-EM structure of FoxP3 bound to T3G repeats uncovered a novel architectural function of FoxP3: it cooperatively forms a multimeric assembly on T3G repeat DNA, where each subunit recognizes the TGTTTGT sequence, in place of the canonical TGTTTAC^14^. This organization enables FoxP3 to create a tightly packed array on DNA, facilitating the bridging of two DNA copies and stabilizing chromatin loops in Tregs^14,26,27^. Supporting this architectural role, FoxP3 is highly expressed in Tregs, with its mRNA level equivalent to approximately 10-30% of those of abundant proteins, such as ribosomal proteins (Figure S1A).

While the structure of FoxP3 bound to T3G repeats provided key insights into how FoxP3 recognizes microsatellites, it also raised many new questions. Chief among them is how this DNA-scaffolded multimeric architecture adapts when the underlying DNA sequence changes. This is important because other TnG repeats, such as T2G, T4G, and T5G, are similarly enriched at FoxP3-occupied sites, with each seen in 20%, 39%, 47% of the high-confidence FoxP3 CUT&RUN (CNR) peaks^27,28^ as compared to 35% for T3G repeats (Figure S1B, S1C). Many FoxP3-bound loci contain a diverse mix of T2G, T3G, T4G and T5G repeats (Figure S1C, iii-v). Furthermore, allelic imbalance analysis for FoxP3’s genomic occupancy using heterozygous mice^28^ showed that these TnG repeats besides T3G repeats can drive FoxP3’s genomic binding (Figure S1D). However, the differences in their repeat unit sizes make the arrangement of FoxP3 on T3G repeats incompatible with these other sequences. In addressing this question, we discovered a remarkable structural plasticity in FoxP3 that allows it not only to recognize diverse TnG repeats but also to expand its multimeric assembly to include 2, 3 or 4 DNA copies within a single complex. Moreover, our study revealed that FoxP3 can leverage and incorporate nucleosomes into its assembly by preferentially targeting nucleosome-mediated local DNA hairpin structures, which in turn facilitate distal DNA bridging and higher-order assemblies. These findings uncover novel modes of TF-DNA interactions and the unexpected role of nucleosome in promoting FoxP3-DNA interactions.

### Structure of FoxP3 multimers in complex with T4G repeat DNA

To explore how FoxP3 binds T4G repeat DNA, we determined the cryo-EM structure of FoxP3 in complex with (T_4_G)_15_ DNA. We used FoxP3 construct without the N-terminal disordered region (FoxP3^ΔN^, Figure 1A upper left), which retains the same DNA sequence specificity as the full-length protein^14,25^. This construct contains the C-terminal forkhead DNA-binding domain (DBD), the hydrophobic loop known as the Runx1-binding region (RBR)^29^, a dimerization coiled-coil domain^30^. Negative stain EM and cryo-EM analyses of the FoxP3–(T_4_G)_15_ DNA complex revealed multimeric architectures (Figure S2A, S2B). Cryo-EM 3D reconstruction identified three distinct classes: two classes with two copies of DNA (DNA^A^ and DNA^B^, and DNA^B^ and DNA^C^) and one with three copies of DNA (DNA^A^, DNA^B^ and DNA^C^) (Figure S2C-2F). The three-DNA structure (Figure 1A) could be reconstituted by aligning the DNA^B^ from the two-DNA structures (Figure S3A). All DNAs displayed a classic B-form double-stranded structure (TnG strand in black, AnC strand in grey in Figure 1). The forkhead DBD formed the canonical winged-helix fold in all cases. Only the DBD and part of the RBR (residue ∼318-336) were traceable, indicating heterogeneous conformations for the rest of the protein.

**Figure 1.**
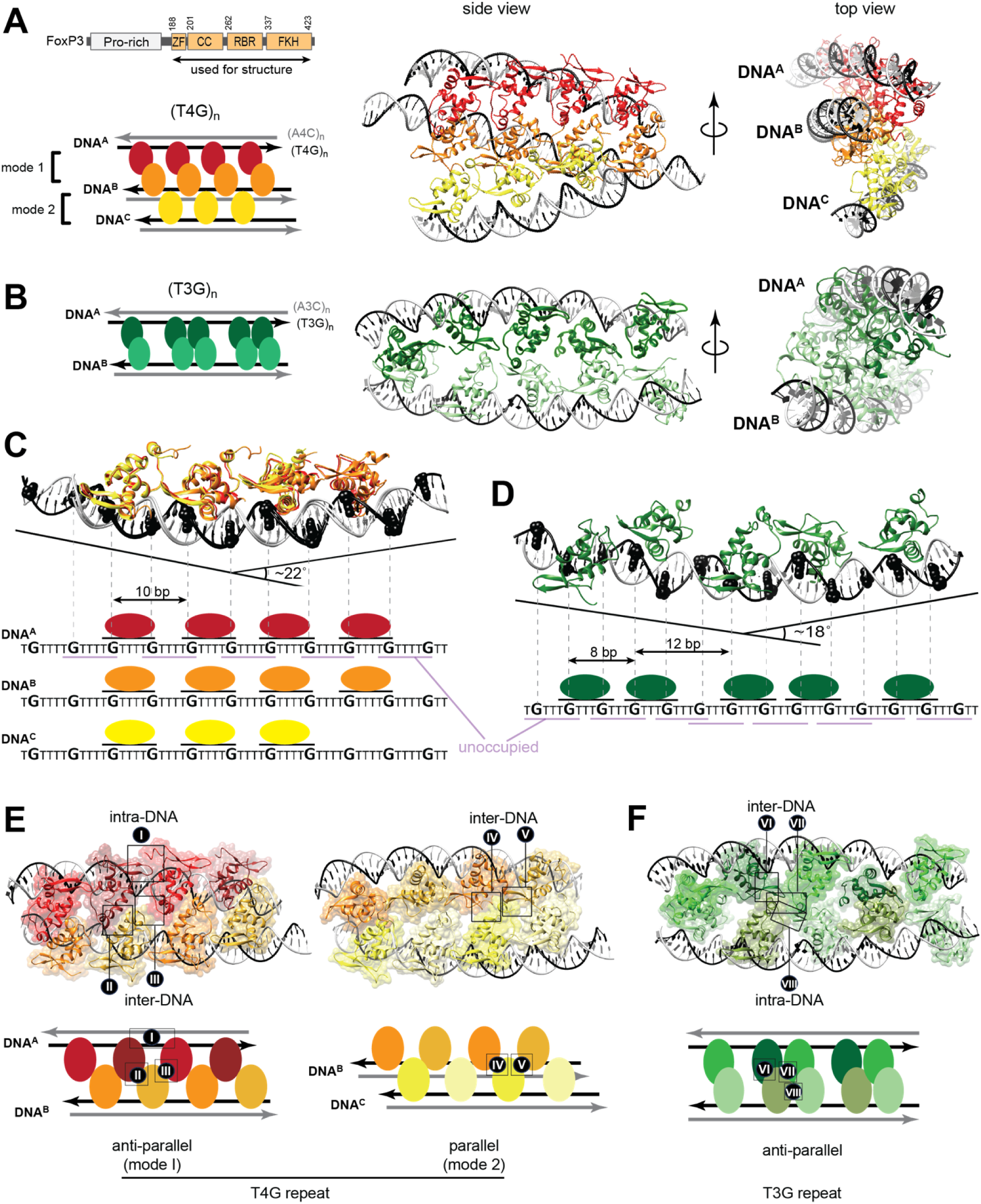
Structure of FoxP3 multimers in complex with T4G repeat DNA. A. Schematic of the domain architecture of FoxP3 (Top left). The N-terminal truncation (FoxP3^ΔN^) was used in all structural and biochemical analyses. Cryo-EM structure of FoxP3^ΔN^ multimer in complex with three DNA molecules harboring (T_4_G)_15_. FoxP3 subunits bound to different dsDNA are colored differently. Within each dsDNA molecule, the T4G repeat strand is in black, whereas the complementary strand is in grey. Schematic (bottom left), side view (middle) and top view (right) are shown. B. Cryo-EM structure of FoxP3^ΔN^ multimer in complex with two DNA molecules harboring (T_3_G)_18_ (PDB:8SRP). Schematic (left), side view (middle) and top view (right) are shown. C. Superposition of FoxP3^ΔN^ subunits bound to T_4_G repeat DNA^A^ (red), DNA^B^ (orange), and DNA^C^ (yellow). Black sphere models in DNA indicate guanosine bases. Individual FoxP3 subunits interact with DNA in the identical manner, recognizing TGTTTTG sequence with a spacing of FoxP3 every 10 bp. This arrangement positions FoxP3 along one side of the DNA, facilitating intra-DNA interactions between adjacent subunits, bending the DNA towards the bound side (∼22°), while leaving behind TGTTTTG sites on the opposite side of the DNA (light purple) unoccupied. D. FoxP3^ΔN^ subunits bound to T_3_G repeat DNA molecules with 18° bent (PDB:8SRP). Each subunit binds DNA recognizing TGTTTGT, leading to alternating spacing of 8 bp and 12 bp. E. Inter-subunit interactions in the FoxP3-T4G repeat complex. One type of intra-DNA interaction (type I) was identified in all subunits. When considering inter-DNA interactions, four types of inter-subunit interactions were identified (type II-V). Type II and III interactions were found in the anti-parallel DNA bridging mode (mode 1), whereas type IV and V in the parallel DNA bridging mode (mode 2). FoxP3 subunits on DNA^A^, DNA^B^, and DNA^C^ are shown in shades of red, orange and yellow, respectively. Unlike in (A) and (C), different colors were used to distinguish adjacent subunits on the same DNA. F. Two inter-DNA interfaces (VI and VII) and one intra-DNA interface (VIII) in FoxP3^ΔN^– (T_3_G)_18_ complex structure (PDB:8RSP).

In all three complex structures, each T4G DNA was directly bound by three to four copies of FoxP3 (Figure 1A, 1C). The FoxP3 molecules bound to different DNAs then interacted with each other, linking DNA^A^ to DNA^B^, and DNA^B^ to DNA^C^ (Figure 1A, middle). A top-down view along the DNA^B^ axis revealed that the three DNA molecules were skewed relative to one another, with proteins predominantly clustered on one side, leading to an asymmetric distribution of proteins around the DNA bundle (Figure 1A, right). These structures, whether involving two or three DNA molecules, were markedly different from the FoxP3 multimers on T3G repeats^14^, which displayed a symmetric bridging of two DNA copies (Figure 1B).

To investigate the differences between the T3G and T4G complexes, we analyzed how FoxP3 subunits interacted with each DNA sequence. Superposition of DNA^A^, DNA^B^ to DNA^C^ of T4G repeats revealed that all FoxP3 subunits bound to each DNA in an identical manner, recognizing every other TGTTTTG sequence within T4G repeats and spacing themselves every 10 base pairs (Figure 1C). This arrangement allowed FoxP3 molecules to align on one side of the DNA, forming direct protein-protein interactions with adjacent subunits (type I, intra-DNA interaction in Figure 1E) and inducing a ∼20° inward bend in the DNA (Figure 1C). This DNA bending likely prevented FoxP3 from binding the unoccupied TGTTTTG sites on the opposite side of the DNA by precluding intra-DNA interactions (Figure S3B). When binding to T3G repeats, FoxP3 also caused inward bending^14^ (Figure 1D); however, it recognized the TGTTTGT sequence with alternating spacings of 8 and 12 base pairs. In this case, intra-DNA interactions occurred only at the 8 base pair spacing, not at 12 base pairs (type VIII interaction in Figure 1F).

The DNA-bridging mechanisms also differed between T3G and T4G repeats. With T4G repeats, DNA^A^ and DNA^B^ were antiparallel in sequence orientation, whereas DNA^B^ and DNA^C^ were parallel, showing at least two distinct ways FoxP3 can bridge T4G DNA. In both cases, FoxP3 proteins interdigitated between the two bridging DNA molecules, with each protein on one DNA forming interactions with two proteins on the other DNA (inter-DNA interactions II-V in Figure 1E, further discussed in Figure 3). In contrast, only one mode of bridging (antiparallel) was observed with T3G DNA. Even when comparing with the antiparallel inter-DNA interaction on T4G DNA, the T3G complex showed different protein-protein interactions (type VI, VII, Figure 1F).

Thus, a slight change in microsatellite repeat unit from T3G to T4G can drastically alter the multimeric assembly architecture by modifying inter-subunit spacing and interactions. The T4G structures also demonstrate that even with a single T4G repeat sequence, DNA bridging interactions can occur in at least two distinct modes––antiparallel (mode 1) and parallel (mode 2) using distinct interfaces. Moreover, these complexes can expand from two-DNA to three-DNA complexes by combining different modes of DNA bridging.

### Structure of FoxP3 multimers in complex with T2G repeat DNA

We next determined the cryo-EM structure of FoxP3 in complex with (T_2_G)_24_ DNA. Both negative stain EM and cryo-EM images suggested larger particles of T2G complexes, compared to T3G and T4G complexes (Figure S4A, S4B). This was consistent with the slower migration rate of the T2G complex by EMSA (Figure S5A) and earlier elution by size-exclusion chromatography (Figure S5B). Cryo-EM reconstruction revealed two classes of the FoxP3–T2G complex: a major class (93%) containing ∼28 copies of FoxP3 and 4 copies of DNA, and a minor class (7%) that was half the size (Figure S4C). The major complex displayed a barrel-like structure where the four DNAs (DNA^A^-DNA^D^) formed the vertical pillars, each bound by 7 copies of FoxP3 (Figure 2A, Figure S4C-4F). The FoxP3 subunits formed interfaces between adjacent DNA molecules, connecting DNA^A^ with DNA^B^, DNA^B^ with DNA^C^, DNA^C^ with DNA^D^, and DNA^D^ with DNA^A^ (Figure 2A).

**Figure 2.**
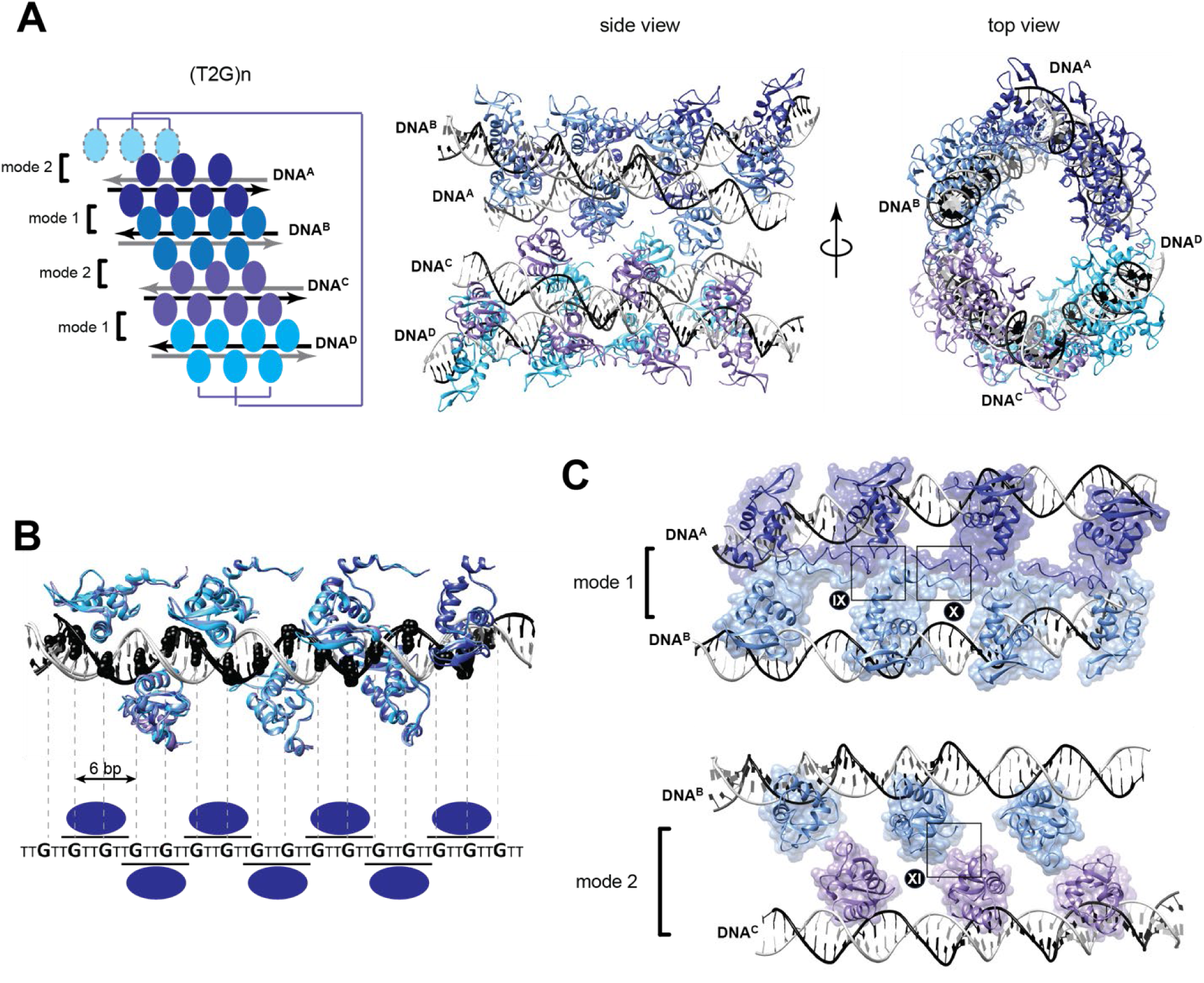
Structure of FoxP3 multimers in complex with T2G repeat DNA. A. Cryo-EM structure of FoxP3^ΔN^ multimer in complex with four DNA molecules harboring (T_2_G)_24_. Within each dsDNA molecule, the T4G repeat strand is in black, whereas the complementary strand is in grey. FoxP3 subunits bound to different dsDNA are colored differently. Schematic (left), side view (middle) and top view (right) are shown. B. Superposition of FoxP3^ΔN^ subunits bound to T2G repeat DNA^A^ (navy blue), DNA^B^ (blue), DNA^C^ (purple) and DNA^D^ (sky blue). Guanosine bases are indicated with spheres. Individual FoxP3 subunits interact with DNA in the identical manner, recognizing TGTTGTT sequence with a spacing of FoxP3 every 6 bp. This leads to FoxP3 molecules on each DNA forming two linear arrays on opposite sides of the DNA. C. Inter-subunit interactions in the FoxP3-T2G repeat complex. Three distinct inter-DNA interactions (type IX-XI) were observed. Type IX and X interactions were found in mode 1 bridging, whereas type XI in mode 2 bridging. Both modes are in the antiparallel orientation. No clear density for intra-DNA interaction was identified.

Superposition of the four T2G DNA molecules revealed that all FoxP3 subunits bound to each DNA in an identical manner, recognizing every other TGTTGTT sequence within T2G repeats and spacing themselves every 6 base pairs along the DNA (Figure 2B). This arrangement placed adjacent FoxP3 subunits on opposite sides of the DNA, resulting in no visible intra-DNA contact between FoxP3 proteins. As a consequence, T2G repeat DNA showed little bending as seen with the T3G or T4G repeats (Figure 2B).

To compensate for the lack of intra-DNA interactions, there were extensive inter-DNA interactions mediated by FoxP3 proteins. Two modes of DNA bridging were observed: mode 1 between DNA^A^ and DNA^B^ and between DNA^C^ and DNA^D^, and mode 2 between DNA^B^ and DNA^C^ and between DNA^D^ and DNA^A^ (Figures 2A, 2C). While both modes of bridging were between two DNAs in antiparallel orientations, the nature of the protein-protein interactions involved were distinct in mode 1 and 2. Mode 1 bridging involved eight copies of FoxP3 (four on each DNA) with 2 types of inter-subunit interactions (type IX and X, Figure 2C), exhibiting strong density indicative of ordered interactions. In contrast, mode 2 bridging involved six copies of FoxP3 (three on each DNA, type XI in Figure 2C) and showed weaker density (Figure S4G), suggesting weak and heterogeneous interactions.

Supporting the notion that the mode 1 bridging is more efficient, the minor population (7%) of reconstructed particles displayed only the mode 1 interaction (Figure S4C). These findings suggest that the formation of four-DNA complexes may proceed stepwise, beginning with the mode 1, followed by the mode 2 interaction. Importantly, all three interaction types (IX, X and XI) seen in T2G complexes were distinct from one another and differed from those observed in the T3G and T4G complexes. Thus, these results once again demonstrate how small differences in microsatellite repeat size can transform the assembly architectures and inter-subunit interactions.

### FoxP3 shapeshifts to form different assembly architectures

We next compared the FoxP3–FoxP3 interactions within the FoxP3 multimers on T2G, T3G^14^, and T4G repeat DNA, as well as the FoxP3 dimer found on IR-FKHM^25^ (Figure 3A). A total of twelve distinct pairwise interactions were observed across these structures (types I–XI in the FoxP3 multimers and type XII in the dimer). For each interaction, we aligned one subunit at the center and mapped the corresponding partner subunit around it. Symmetrical interactions (denoted with * in Figure 3A) positioned the partner subunit identically, regardless of which molecule was centered, whereas asymmetric interactions (without *) appeared differently depending on which subunit was at the center. This analysis revealed that FoxP3 employs a remarkably diverse and expansive surface area for inter-subunit interactions. Notably, some surfaces were involved in forming two distinct interactions (*e.g.*, types I and IV in the T4G complex), underscoring the versatility of FoxP3’s surface in facilitating homotypic interactions.

**Figure 3.**
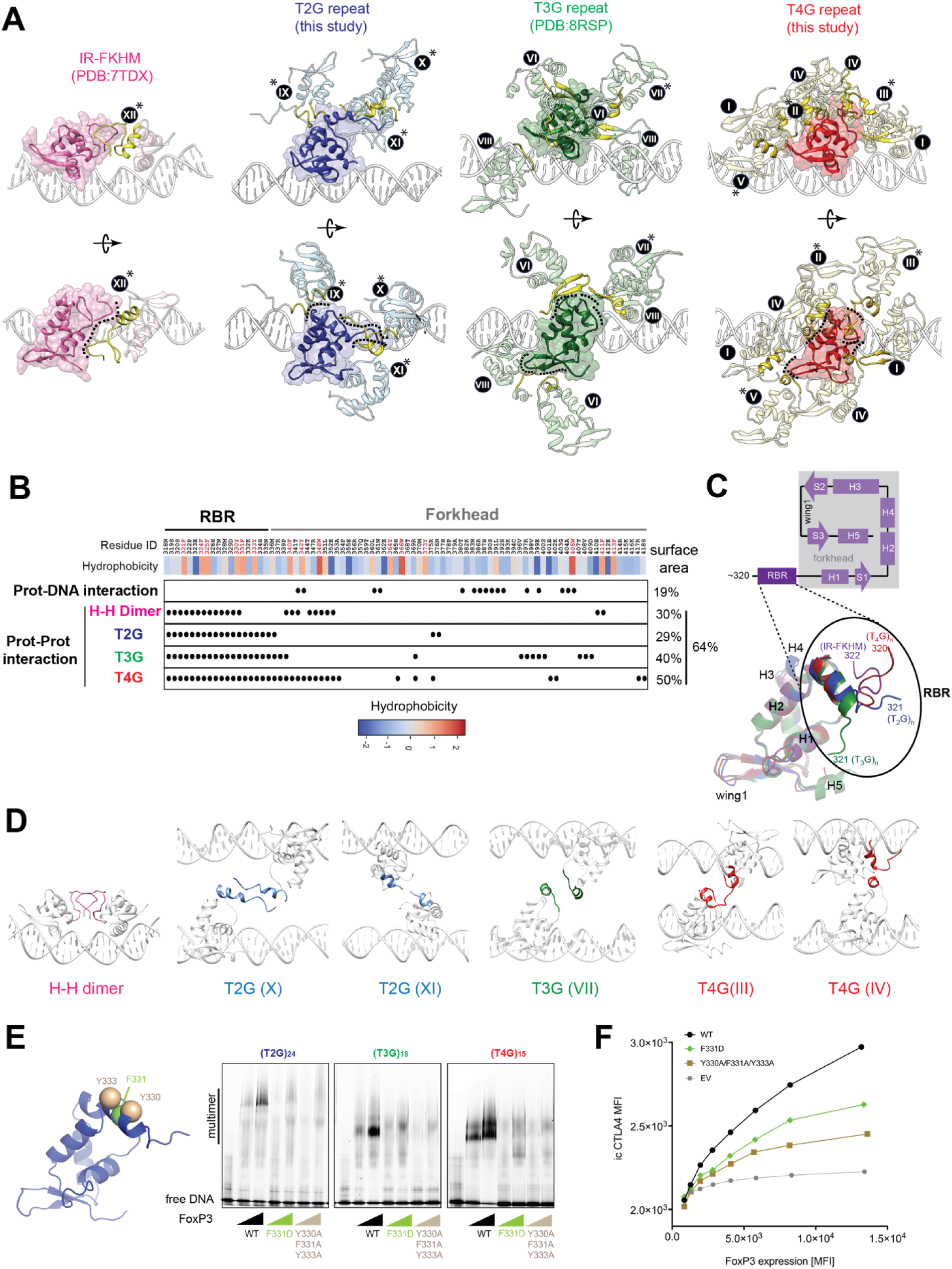
Shapeshifting RBR domain confers structural plasticity to FoxP3 assemblies. A. FoxP3–FoxP3 interactions within the multimeric assemblies of FoxP3 bound to IR-FKHM, T2G, T3G, and T4G repeats DNA. Each pair of interacting subunits was isolated from the complex and centered on one subunit, with the partner subunit mapped around it. Symmetrical pairs (*) exhibit the same interaction mode regardless of which subunit is centered, while nonsymmetrical pairs display two different positions based on the centered subunit. Various types of interactions are superimposed on one subunit and are shown from two perspectives. B. Residues involved in protein-DNA or protein-protein interactions, and their hydrophobicity (black oval). All structures showed the identical protein-DNA interactions. However, protein-protein interactions involved different residues depending on the bound DNA, with the exception of the RBR, which consistently mediates protein-protein interactions across all DNA complexes. Hdrophobicity values were determined using the Wimley–White scale. Aromatic residues were colored red in Residue ID. Only residues with a solvent accessible surface (SAS) >20 Å^2^ in the monomeric state (as calculated by Chimera) are listed. The total surface area involved in protein-DNA or protein-protein interactions was calculated by summing the SAS of all interacting residues in each structure. C. Top: Secondary structure of FoxP3 RBR-forkhead. H indicates α-helix, S indicates β-strand. Bottom: Superposition of representative FoxP3^ΔN^ subunits from the structure in complex with IR-FKHM (purple, PDB: 7TDX), T3G repeat (green, PDB:8RSP), T2G repeat (blue, this study) and T4G repeat (red, this study). D. Six different types of RBR-RBR interaction modes (III, IV, VII, X-XII, defined in Figure 3A) observed in four complex structures. RBRs are indicated with different colors according to the bound DNA sequence. E. Effect of the RBR-RBR interface mutations (F331D or Y330A/F331A/Y333A) on FoxP3 multimerization on T2G, T3G and T4G repeats. FoxP3^ΔN^ (0, 0.8, 1.6 μM) with and without mutations were incubated with 0.2 μM TnG repeat DNA. Sybrgold stain was used for visualization. Left: mutated residues were shown on one of the FoxP3 subunits bound to T2G repeats from the same view as in (C). F. Transcriptional activity of FoxP3 variants. CD4+ T cells were retrovirally transduced to express FoxP3, and its transcriptional activity was analyzed by measuring the protein levels of the known target genes CTLA4 using fluorescence-activated cell sorting (FACS). FoxP3 levels were measured on the basis of Thy1.1 expression, which is under the control of IRES, encoded by the bicistronic FoxP3 mRNA. MFI, mean fluorescence intensity.

To systematically compare the inter-subunit interactions in different FoxP3–DNA complexes, we identified and tabulated residues involved in inter-subunit interactions in each complex (Figure 3B). In all cases, residues involved in DNA interaction were identical. However, residues involved in protein-protein interactions were different. Notably, the T4G complex utilized the most residues, with 50% of the protein surface (within residues 318-414) engaged in protein-protein interactions, followed by 40% for the T3G complex, and 29-30% for the T2G complex and dimer. Altogether, 64% of the surface in the RBR-forkhead region participated in protein-protein interactions, compared to only 19% involved in DNA binding (Figure 3B). These findings suggest that the majority of the FoxP3 surface in this region is primarily dedicated to multimerization rather than DNA binding.

Another notable finding was that the RBR consistently played a central role in all four complexes (Figure 3B). To understand how RBR accommodates such distinct inter-subunit interactions across different complexes, we superimposed a representative subunit from each structure by aligning the forkhead DBD. This analysis showed that RBR adopts different conformations in each structure (Figure 3C). Even when interacting with another RBR, its conformation varied depending on the spacing and orientation of the partner subunit (Figure 3D). This remarkable shape-shifting capability is likely driven by RBR’s unique properties—namely, its high hydrophobicity and the unusually high density of surface-exposed aromatic residues. Of the eighteen traceable RBR residues (residues 318-336), six were aromatic and well-conserved (Figures 5B, S5C), a significantly higher frequency compared to other regions (e.g., 9 out of 81 aromatic residues in the forkhead domain). This hydrophobicity likely enables RBR to fold into diverse structures and interact with different surfaces. Supporting the functional importance of this hydrophobicity, mutations in key aromatic residues (e.g., Tyr330, Phe331, Tyr333), either collectively to alanine or as a single point mutation to aspartic acid, impaired FoxP3’s multimerization on all three DNA sequences (T2G, T3G, and T4G; Figure 3E) and disrupted its cellular functions in CD4 T cell transduction experiments (Figure 3F).

In summary, RBR is involved in all FoxP3 assemblies examined to date, regardless of the DNA sequence or overall shape of the multimeric architecture. This universal role is attributed to RBR’s intrinsic shapeshifting ability, which allows it to adopt different multimeric assemblies, a capability facilitated by its hydrophobic nature.

### FoxP3 multimers exhibit ultrastability across diverse DNA copy numbers

The observed heterogeneity in the copy number of DNA across our FoxP3 multimeric structures raised a question about whether there are many additional assembly states of FoxP3 complexes with TnG repeats that were not captured in the structures and how stable each assembly structure is. For example, the four-DNA complex was the predominant species (93%) of FoxP3 bound to T2G repeats, with the two-DNA complex being a minor species (7%). This raises the question how stable the two-DNA complex is and whether other intermediate complexes with varying DNA copy numbers could form and persist under physiological conditions.

To address these questions, we employed single-molecule analysis, starting with testing whether FoxP3 can stably bridge two copies of T2G, T3G, and T4G repeats using a “nunchuck” construct labeled for single molecule fluorescence resonance energy transfer^31,32^ (Figure 4A, left). In this construct, two dsDNA regions with identical TnG repeat sequences––(T_2_G)_24_, (T_3_G)_18_, (T_4_G)_15_–– were connected by an 18-atom hexa-ethyleneglycol spacer (TnG-TnG, Figure 4A, left). The two dsDNA regions were labeled with Cy3 (FRET donor) and Cy5 (FRET acceptor) at their centers. The nunchuck was immobilized on a streptavidin-coated surface and extensively washed to ensure elimination of free nunchucks.

**Figure 4.**
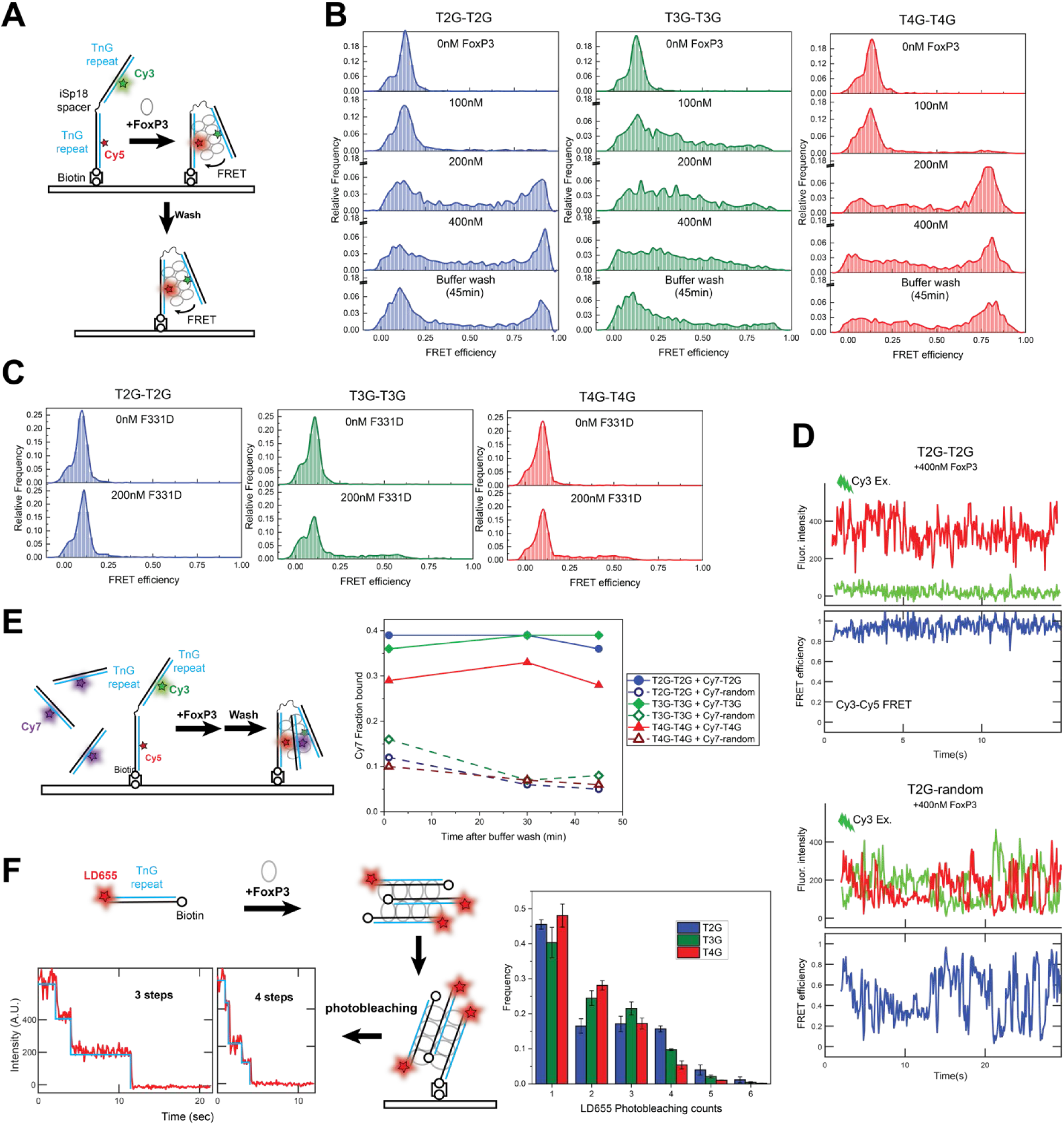
FoxP3 multimers exhibit ultrastability across diverse DNA copy numbers. A. TnG-TnG “nunchuck” DNA construct used in the single molecule analysis. The construct consists of two dsDNA regions with identical TnG repeat sequences––(T_2_G)_24_, (T_3_G)_18_, (T_4_G)_15_––were connected by an 18-atom hexa-ethyleneglycol spacer. Each TnG repeat is labeled with Cy3 or Cy5. FoxP3-induced DNA bridging between the Cy3 and Cy5-labeled repeats was measured by Cy3-Cy5 FRET. B. Distributions of FRET efficiencies for TnG-TnG with an increasing concentration of FoxP3. Stability of bridged DNA complex was measured by washing with buffer lacking FoxP3 for 45 min. C. Effect of F331D mutation on FRET for different TnG repeats. D. Representative time traces for fluorescent intensity and Cy3-Cy5 FRET efficiency of T2G-T2G (top) and T2G-random (bottom) nunchucks, both in the presence of 400 nM FoxP3. E. Higher-order multimerization of FoxP3 on immobilized TnG-TnG nunchucks in the presence of free Cy7-labeled DNA. Cy7-DNA, containing either TnG repeats or random sequences, was added to immobilized nunchucks along with FoxP3 (300 nM), and were washed with buffer lacking FoxP3 for the indicated times. FoxP3-induced FRET (Figure S8A) and recruitment of Cy7-DNA (Figure S8B) were measured. The fraction of bound Cy7-DNAs was plotted against washing time. F. Single-molecule pulldown of LD655-labeled TnG repeat DNA incubated with FoxP3 (1 μM) in solution prior to surface immobilization. Number of LD655-DNA within the FoxP3 complex was measured as a function of LD655 photobleaching counts. Right: distribution of the photobleaching counts for different TnG repeats.

In the absence of FoxP3, the nunchuck constructs exhibited low FRET, indicating an extended conformation (Figure 4B). However, upon the addition of FoxP3 (100-200 nM), FRET increased for all three TnG repeats, suggesting folding of the nunchuck construct (Figure 4B). The FRET signal remained stable even after washing with buffer devoid of FoxP3 for 45 minutes (Figure 4B). This indicates the ultrastability of the multimeric complexes once they are formed. Notably, FRET increase was minimal when FoxP3 was mutated (F331D) to impair multimerization (Figure 4C), or when one arm of the nunchuck was mutated to a random sequence (TnG-random, Figure S6A). At higher FoxP3 concentrations (300-400 nM), some FRET was observed with the TnG-random sequence (Figure S6A), but these signals were unstable (Figure 4D, Figure S6B) and rapidly lost within 1 min of washing (Figure S6A). These results demonstrate that FoxP3 forms ultrastable two-DNA complexes with all three TnG (n=2-4) repeats, but not with the random sequence.

We noticed that the T3G-T3G nunchuck behaved differently from T2G-T2G and T4G-T4G nunchucks. The T3G-T3G nunchuck was more sensitive to FoxP3, requiring a lower concentration (100 nM) of FoxP3 to induce FRET, compared to 200 nM for T2G-T2G and T4G-T4G (Figure 4B). Between T2G-T2G and T4G-T4G, a higher fraction of T4G-T4G showed an increase in FRET upon FoxP3 addition, suggesting that T2G repeats is the least efficient among the three sequences in forming a two-DNA complex, consistent with our structural findings. Additionally, T3G-T3G exhibited a broader FRET distribution, indicating variable Cy3-Cy5 distances when folded by FoxP3 (Figure 4B). This contrasts with the bimodal distribution seen with T2G-T2G and T4G-T4G, which indicates minimal Cy3-Cy5 distance when folded by FoxP3 (Figure 4B). These differences suggest that FoxP3 binds and bridges T3G repeats more efficiently, tolerating shorter overlaps between bridged T3G DNAs, whereas T2G and T4G may require maximal DNA overlap to achieve stable binding. This is consistent with the previous observation that FoxP3 has a higher affinity for T3G repeats than for T2G or T4G repeats^14^.

Next, we examined FoxP3’s capacity to bridge more than two DNA copies for T2G, T3G, and T4G repeats by monitoring the ability of the folded nunchucks to recruit separate, Cy7-labeled DNA sharing the same repeat sequence as the nunchuck (TnG^Cy7^, Figure 4E, left). Upon addition of FoxP3 (300 nM), Cy3-Cy5 FRET again increased and the FRET signal was stable against buffer washing, as was in the absence of TnG^Cy7^ (Figure S7A). Approximately 30-40% of the nunchucks recruited TnG^Cy7^ and stably retained TnG^Cy7^ against 45 min of washing (Figure 4E, Figure S7B). Minimal TnG^Cy7^ recruitment was observed in the absence of FoxP3 (Figure S7B) or when Cy7-labeled DNA had a random sequence (Figure 4E, right). Photobleaching analysis revealed that most Cy7-positive complexes contained one or two TnG^Cy7^, corresponding to three- or four-way DNA bridging (Figure S7C).

To further quantify DNA copy numbers within the FoxP3 multimers, we conducted a single-molecule pull-down assay^33^, where FoxP3 was incubated with TnG repeat DNA in solution prior to surface immobilization (Figure 4F). DNA was pre-labeled with the photostable fluorophore LD655, allowing accurate measurement of DNA copy number by photobleaching (Figure 4F). About 50-60% of the spots contained two or more DNA molecules, with a significant fraction containing three or four (Figure 4F). T2G repeats had the highest propensity for forming four-DNA complexes and the lowest for two-DNA complexes, consistent with other biochemical and structural observations (Figure 2, Figure S5).

Overall, these single-molecule analyses demonstrate the remarkable heterogeneity and stability of FoxP3 multimers formed with varying copy numbers of TnG repeat DNA.

### Nucleosomes further enhance FoxP3’s ability to bridge T_n_G repeats

Our single-molecule analyses suggested that FoxP3-mediated DNA bridging can occur both locally (in cis) and distally (in trans), and that DNA bending could play a role in local DNA bridging. We thus asked whether DNA-bending proteins, such as nucleosomes, can facilitate FoxP3 to induce local bridging of nearby TnG repeats.

Analyses of TnG repeats within FoxP3 CNR peaks^27,28^ revealed two key features supporting this hypothesis. First, TnG repeats were more densely clustered in regions occupied by FoxP3 compared to those in FoxP3-depleted areas (Figure 5A). These TnG repeats appeared as either long contiguous stretches or multiple discontinuous patches (Figure S1C), which could facilitate FoxP3 binding and local folding. Second, while FoxP3 is known to occupy accessible, open chromatin regions (OCRs), FoxP3-occupied TnG repeats often localized near the edges of OCRs, as measured by ATAC-seq^34^ (Figure S8A). This was further confirmed by measuring the distance of TnG repeat centers to the closest ATAC peak summits; these distances were greater than those for other TF motifs (such as ETS and CTCF) known to localize near OCR centers^35^ (Figure 5B). Consistent with this observation, FoxP3-occupied TnG repeats displayed lower accessibility (lower ATAC-seq intensity) than ATAC-seq summits (Figure 5C, S8B), although they were significantly more accessible than TnG repeats outside FoxP3 CNR peaks. Additionally, TnG repeats within FoxP3 CNR peaks exhibited lower accessibility than ETS or CTCF motifs (Figure 5C, S8B). Given that nucleosomes are key factors limiting chromatin accessibility and that TFs at OCR edges often interact with histones^35^, we hypothesized that nucleosomes might play a role in FoxP3 binding.

**Figure 5.**
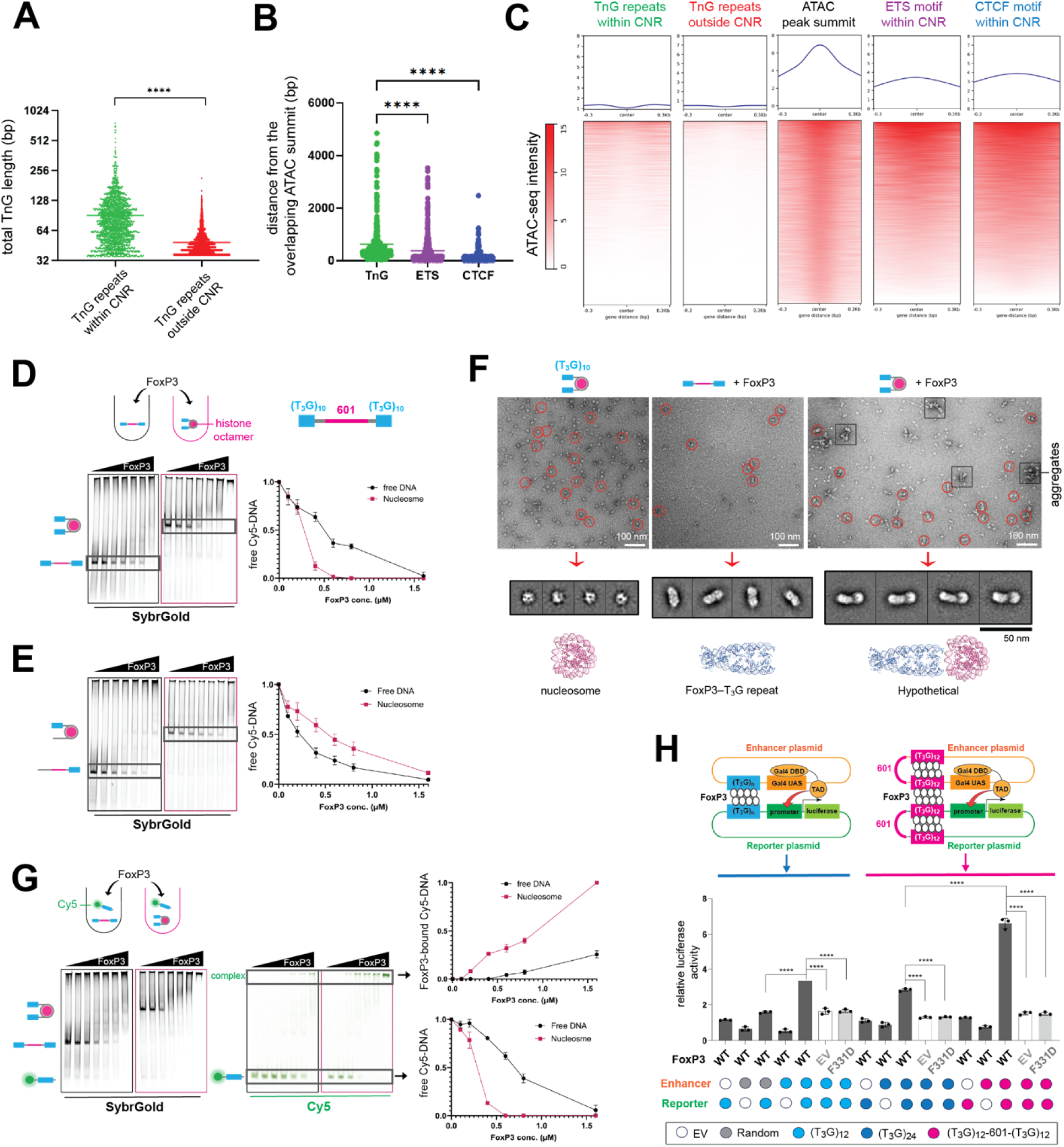
Nucleosomes can facilitate FoxP3 bridging of T3G repeats. A. Total length of TnG repeats (in basepairs) within 1 kb window centering at the TnG repeat sites within (n=2,019) or outside FoxP3 CNR peaks (n=3,579). For TnG repeats outside FoxP3 CNR, only the sites within Treg ATAC peaks that are at least 10 kb away from the FoxP3 CNR peaks were considered. FoxP3 CNRs are the union of the Rudensky and Dixon-Zheng peaks as defined in Figure S1B. Statistical analysis was performed using two-tailed unpaired *t*-tests; \*\*\*\**P* < 0.0001. B. Distance from ATAC summits to the closest center of TnG region, ETS motif or CTCF motif within Foxp3 CNR peaks. Statistical analysis was performed using one-way ANOVA; \*\*\*\**P* < 0.0001. Previously reported thymic Treg ATAC-seq data (PMID:27992401) were used. C. Comparison of ATAC-seq intensity within 600 bp window, centered on TnG repeats within or outside Foxp3 CNR peaks, summits of ATAC peaks overlapping with Foxp3 CNR peaks, ETS or CTCF motifs within Foxp3 CNR peaks. D. FoxP3 binding to free DNA and nucleosomal DNA harboring (T_3_G)_10_-601-(T_3_G)_10_ as measured by native gel-shift assay. Free DNA and nucleosome (50 nM each) were incubated with an increasing concentration of FoxP3 (0, 0.1,0.2,0.4, 0.6, 0.8, 1.6 μM). Sybrgold stain was used for visualization and quantitation for FoxP3-free DNA species. E. FoxP3 binding to free DNA and nucleosomal DNA harboring (T_3_G)_10_-601-random sequence as measured by native gel-shift assay. Experiments were performed as (A). F. Top: Representative negative stain electron micrographs of nucleosomal DNA without FoxP3 (left), nucleosome-free DNA with FoxP3 (middle), and nucleosomal DNA with FoxP3 (right). The same DNA harboring (T_3_G)_10_-601-(T_3_G)_10_ was used in all cases. Aggregate particles, more pronounced in nucleosomal DNA with FoxP3, were shown in black boxes. Particles in red circle were picked for further 2D classification (middle). Bottom: structures of nucleosome (PDB:7OHC) and FoxP3-bound T3G repeat (PDB:8RSP) and a hypothetical model of nucleosomal (T_3_G)_10_-601-(T_3_G)_10_ DNA bound by FoxP3. G. Trans effect of nucleosome on FoxP3’s interaction with Cy5-labeled (T_3_G)_15_ DNA. 50 nM of free or nucleosomal DNA harboring (T_3_G)_10_-601-(T_3_G)_10_ were incubated with an increasing concentration of FoxP3 (0, 0.1,0.2,0.4, 0.6, 0.8, 1.6 μM) firstly, and then with 20 nM of Cy5 labeled (T_3_G)_15_ DNA prior to gel analysis. Sybrgold stain (left) and Cy5 fluorescence (middle) were used for visualization. Cy5 intensity of FoxP3-free species was quantitated (right). H. Top: Dual luciferase assay schematic. Two types of plasmids were generated: an enhancer plasmid containing an enhancer element (UAS) which can be bound by Gal4 DBD, and reporter plasmid containing the firefly (FF) luciferase gene driven by a minimal promoter. Both plasmids contained (T_3_G)n repeats (n=12 or 24) or (T_3_G)_12_-601-(T_3_G)_12_, upstream of the promoter or enhancer. These two plasmids, along with the FoxP3-expressing plasmid and Renilla luciferase-encoding transfection control plasmid, were co-transfected into 293T cells expressing Gal4 DBD fused with a transcriptional activation domain (TAD) from the unrelated TF Aire (Gal4-TAD). Bottom: Relative level of FF luciferase activities (normalized to Renilla) were shown. EV indicates an empty vector. Random indicates an enhancer or reporter plasmid with T3G repeat replaced by a random sequence of the same length. Data are representative of three biological replicates and presented as mean ± s.d. *p*-values (one-way ANOVA with Tukey’s multiple comparisons test) were calculated for each comparison, **** *p* < 0.0001.

To explore the role of nucleosomes and TnG clustering in FoxP3 multimer assembly, we examined whether a nucleosome positioned between TnG repeats could fold the DNA, bringing the TnG stretches closer together and enhancing FoxP3 binding. Using DNA with a nucleosome-positioning sequence (601) flanked by two stretches of T3G repeats, we found that FoxP3 preferred nucleosomal DNA over free DNA (Figure 5D). Similar preference was observed with T2G and T4G repeats (Figure S9A, S9B). This enhanced binding was not due to direct interaction between FoxP3 and the nucleosome, as FoxP3 binding to DNA with a single TnG repeat, instead of two, was retarded by the nucleosome presence (Figure 5E). Notably, the histone modification H3K27ac, often enriched in FoxP3-occupied genomic sites^36^, did not further enhance FoxP3 binding (Figure S9C). When visualized by negative stain EM, FoxP3 in complex with nucleosomal DNA displayed a stalk-like FoxP3 multimers capped with a round-shaped nucleosome (red circles in Figure 5F for T3G, Figure S9D for T2G, Figure S9E for T4G), which were absent in either the nucleosomal DNA or FoxP3 multimers alone. They were also absent from FoxP3 in complex with nucleosomal DNA harboring a single TnG repeat (Figure S9F). These results support the idea that the nucleosome facilitates FoxP3 binding by bending the DNA and bringing TnG repeats closer.

Negative stain EM also showed that FoxP3 complexes on TnG-TnG DNA exhibited more aggregation in the presence of nucleosomes than without (black boxes in Figure 5F, Figure S9D, Figure S9E). We hypothesized that nucleosomal DNA might act as a platform to recruit not only FoxP3 but also additional TnG DNA *in trans* through multiway bridging. To test this hypothesis, we investigated whether FoxP3 binding to Cy5-labeled TnG DNA was influenced by the presence of separate TnG-TnG DNA either harboring or lacking a nucleosome (Figure 5G). Consistent with our hypothesis, the presence of nucleosomal DNA enhanced FoxP3 binding to a separate Cy5-TnG DNA, as indicated by both a reduction in free Cy5-DNA and an increase in FoxP3-bound Cy5-DNA (Figure 5G). These demonstrate that nucleosome occupancy positively influences FoxP3 multimerization both *in cis* and *in trans*.

To test the role of nucleosomes in FoxP3 multimerization in cells, we developed a reporter assay using two plasmids: a reporter plasmid containing the luciferase gene driven by a minimal promoter, and an enhancer plasmid with an enhancer element (upstream activation sequence, UAS) to be bound by Gal4 DBD fused with a transcriptional activation domain (TAD) from an unrelated TF AIRE^37^. Both plasmids contained T3G repeats (12 or 24 copies), with and without the 601 sequence, upstream of the promoter or enhancer (Figure 5H). Both plasmids, along with a FoxP3-expressing plasmid, were co-transfected into the 293T cell line expressing Gal4DBD-TAD. In the presence of FoxP3, the reporter activity increased when the enhancer plasmid contained (T_3_G)_12_, but not when it contained a random sequence of the same length (Figure 5H, samples 1-5). This enhancer-driven activation was not observed without FoxP3 or when using the FoxP3 multimerization mutant F331D (Figure 5H, samples 5-7). This suggests that FoxP3 induced the reporter activity by multimerizing on both the enhancer and promoter plasmids and bridging them. A similar FoxP3-dependent activation was observed with (T_3_G)_24_ (Figure 5H, samples 8-12). Interestingly, inserting the 601 sequence in the center of the (T_3_G)_24_ sequence (Figure 5H) in both the enhancer and reporter plasmids significantly increased the reporter activity (Figure 5H, samples 13-17). These results suggest that FoxP3-mediated enhancer-promoter bridging is further enhanced by the presence of flanking nucleosomes in cells, consistent with the in vitro findings.

## Discussion

Protein homo-multimerization is a fundamental aspect of biology, yet its role in transcription is only beginning to be understood. Unlike protein-only multimerization, DNA-scaffolded TF multimerization is inherently influenced by the underlying DNA sequence. This raises the question of how a TF like FoxP3 can recognize and adapt to similar yet distinct, evolutionarily variable TnG repeat microsatellites. Our structural studies revealed that FoxP3 achieves this using structural plasticity, forming multiple distinct supramolecular assemblies that adjust with repeat periodicity while maintaining a consistent local DNA-binding mode (Figure 6A). This adaptability is driven in part by the shapeshifting RBR loop, which readily adjusts to different spacings and orientations of partner subunits. Additionally, FoxP3 utilizes the extensive surface area (64%) of its forkhead DNA-binding domain for protein-protein interactions—far exceeding the area used for DNA binding (19%)—enabling at least 12 unique pairwise interactions that can be deployed in various combinations to maximize intersubunit connections (Figure 6A). This flexibility allows FoxP3 to recognize a wide array of TnG repeats, as well as the canonical FKHM, making structural adaptability a key feature of FoxP3’s evolutionary response to the dynamic microsatellite landscape.

**Figure 6.**
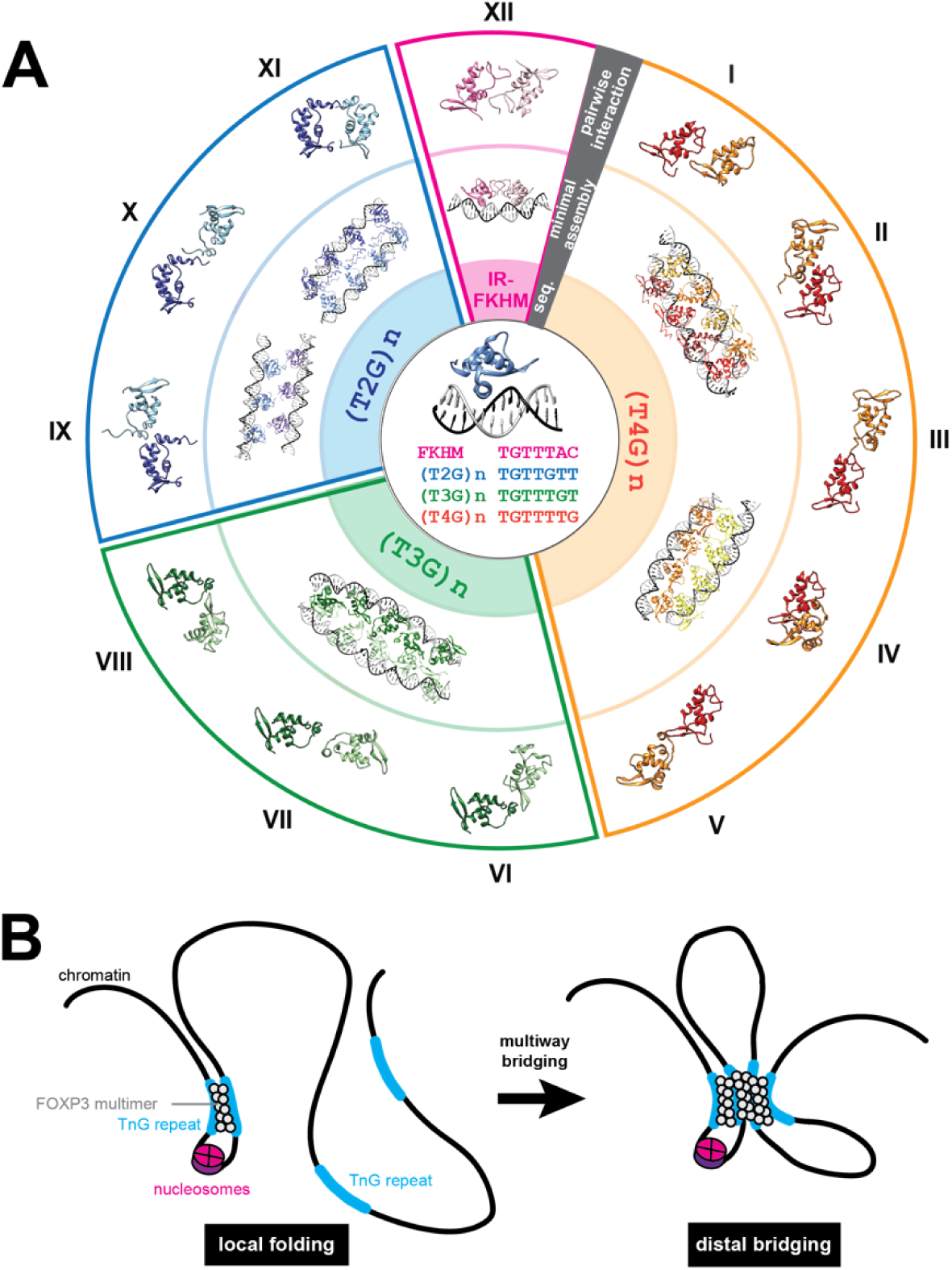
FoxP3 assembly ensembles on microsatellites. A. Diverse FoxP3 assembly states. Regardless of the FoxP3 structure, all assemblies share a DNA-binding mode that recognizes TGTT as the first four nucleotides (center). However, depending on the surrounding DNA sequence (second layer), FoxP3 forms different complexes (third layer). For simplicity, only the minimal complexes are shown— specifically, the H-H dimer on IR-FKHM and a two DNA-bridged structure with TnG repeats. Even a single DNA sequence can induce FoxP3 to form multiple distinct assemblies, which can expand to higher-order structures bridging three to four DNA duplexes (not shown). This architectural complexity is driven by diverse pairwise interactions between FoxP3 subunits (outer layer; 12 identified to date, including 8 discovered here). The presence of mixed TnG repeats in many FoxP3-occupied genomic regions likely supports a range of hybrid assemblies, contributing to substantial structural heterogeneity. B. Hierarchical assembly of FoxP3 multimers through multiway DNA bridging. While FoxP3 can simultaneously bridge three to four distal DNA elements, our data suggests that local DNA folding can significantly facilitate this process. Pre-organized TnG repeat elements, such as those brought together by nucleosomes, can be initially bridged, serving as nucleation sites for recruiting additional DNA elements through FoxP3-mediated multiway bridging. This process leads to dense transcriptional ’hubs’ enriched with regulatory elements, such as enhancers, where FoxP3 localizes.

FoxP3’s architectural versatility extends to multiway DNA bridging—an emerging concept with significant implications for transcriptional regulation but limited mechanistic insights^38–40^. Our data show that FoxP3 can expand its multimeric architecture, bridging not only two DNA duplexes but also three or four. While multiway bridging could occur through simultaneous joining of three or four TnG repeats, our data suggest an alternative, more favorable pathway involving hierarchical assembly. This process likely begins with pairwise bridging of two closely positioned TnG repeats, which then serves as a platform for recruiting distal DNA elements through multiway bridging (Figure 6B). Regardless of the specific mechanism, FoxP3’s capacity to bridge multiple DNA duplexes within a single stable complex supports a model in which it facilitates the formation of dense transcriptional ’hubs’ enriched with regulatory elements like enhancers, which FoxP3 is known to bind^36^. Our results thus add a new layer of mechanistic insight into previous reports that FoxP3 reinforces chromatin loops in Tregs^26,27^.

Finally, our work suggests that FoxP3 can leverage nucleosomes to enhance its DNA target recognition. While FoxP3 is known as a non-pioneering TF that relies on pre-existing chromatin landscapes^15,41,42^, FoxP3-bound TnG repeats often occur near the edges of open chromatin regions (OCRs)—areas typically occupied by pioneering TFs^35^. Our data suggest that this positional specificity may stem from FoxP3’s unique ability to leverage nucleosomes without directly interacting with or displacing them. Nucleosomes enhance FoxP3 assembly by inducing local DNA bending, facilitating local chromatin folding (Figure 6B). Given the ultrastability of FoxP3 multimeric assemblies, we speculate that FoxP3 may reinforce chromatin boundaries and lock in chromatin conformations, functioning as a robust barrier to nucleosome remodeling. These functions may align with recent observations that FoxP3’s role is primarily in amplifying global transcriptomic changes that initiate prior to FoxP3^43^.

In summary, our findings suggest that FoxP3 represents a new class of transcription factor, one capable of adopting multiple distinct conformations to accommodate variable microsatellite sequences, while providing structural stability that can reinforce chromatin boundaries, stabilize nucleosome positioning, and integrate local and distal genomic folding.

## SUPPLEMENTARY FIGURES

**Figure S1.**
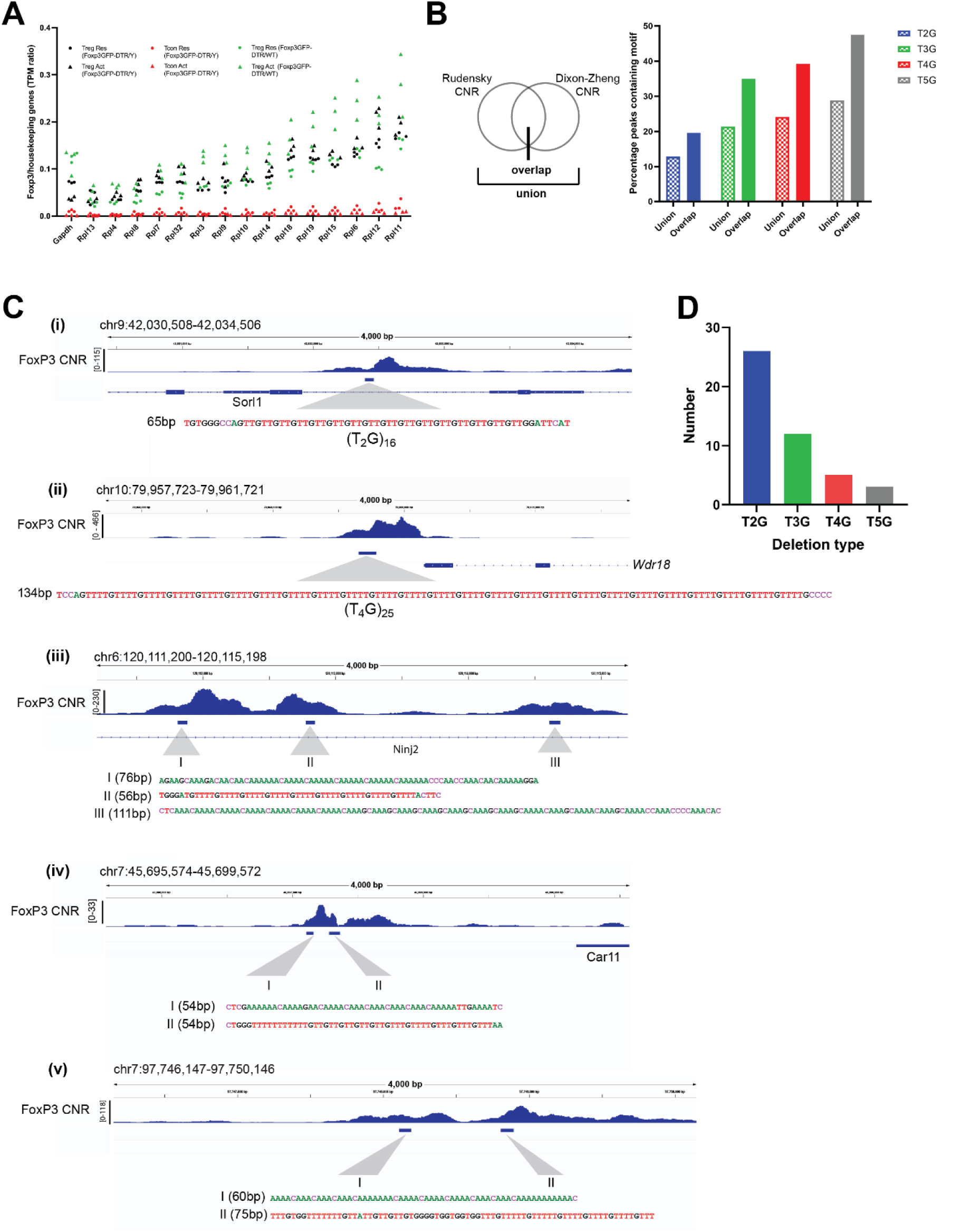
Analysis of FoxP3-occupied T_n_G repeats in the genome. A. Analysis of Foxp3 expression levels in Treg and Tcon cells, at both resting and activated states based on previously published RNA-seq (PMID:17136045, 17136045). X axis: housekeeping genes for normalizing FoxP3 level. Y axis: TPM ratio of FoxP3 over different housekeeping genes in T cells from Foxp3GFP-DTR/Y (male mice expressing human DTR and EGFP genes from the Foxp3 locus, without disrupting expression of the endogenous Foxp3 gene, PMID:17136045) or Foxp3GFP-DTR/WT (heterozygous female mice with normal endogenous Foxp3 expression, PMID:17136045). B. Left: FoxP3 CNR union (n=9,062) and overlap (n=1,354) peaks were derived from the comparison of the Rudensky (PMID:33176163) and Dixon-Zheng datasets (PMID:37932264). Right: percentage of Foxp3 CNR union and overlap peaks containing T2G, T3G, T4G and T5G motifs. C. Genome browser views of Foxp3 CNR peaks harboring T2G and T4G repeats. Some peaks contain clean T2G or T4G repeats (i and ii), while many others contain a mixture of T2G, T3G, T4G and T5G repeats (iii-v). Loci (iii-v) also show clusters of two or more stretches of TnG repeats within 4 kb. Tracks (upper to lower): Foxp3 CNR, TnG regions within Foxp3 CNR peaks, and Refseq gene annotation. D. Number of FoxP3 CNR peaks harboring TnG repeats where deletion in one allele is accompanied by reduction in FoxP3 CNR intensities from the previously published B6/cast F1 hybrid CNR data (PMID:33176163). Among the top 200 sites showing significant B6 or cast bias, 47 of which show TnG deletion in the weaker allele.

**Figure S2.**
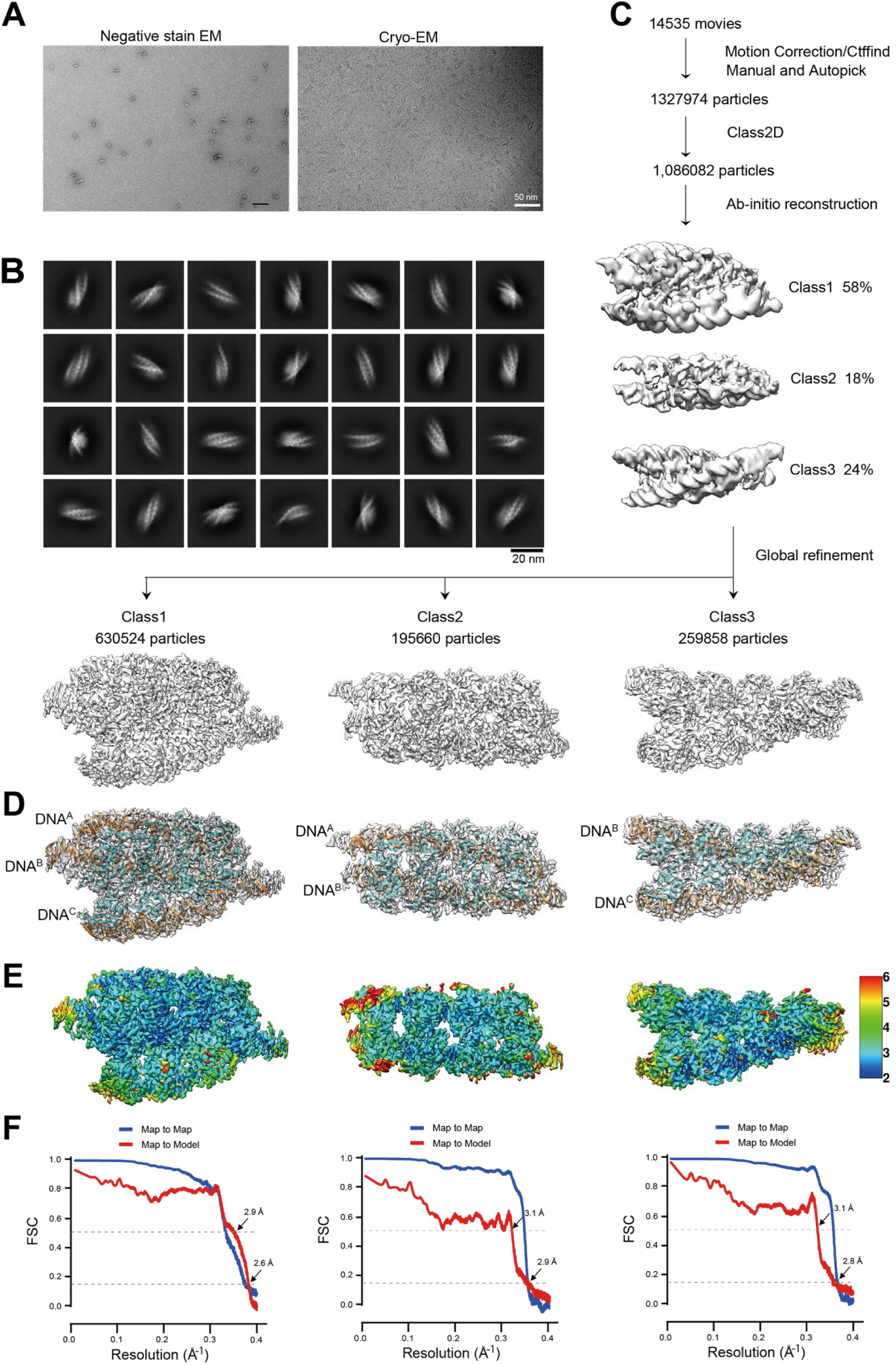
Cryo-EM process of the FoxP3^ΔN^–(T_4_G)_15_ complex. A. Representative negative-stain EM (left) and cryo-EM images (right) of FoxP3^ΔN^ multimers on (T_4_G)_15_ DNA. B. 2D classes chosen for 3D reconstruction. C. Cryo-EM image processing workflow. See details in Methods. D. Cryo-EM maps and ribbon models of FoxP3^ΔN^ multimer in complex with two or three copies of (T_4_G)_15_ DNAs. DNA molecules are colored orange. FoxP3^ΔN^ subunits are colored sky blue. E. Local resolution of Cryo-EM maps was calculated by CryoSPARC. Resolution range was indicated according to the color bar. F. Fourier shell correlation (FSC) curve. Map-to-Map FSC curve was calculated between the two independently refined half-maps after masking (blue line), and the overall resolution was determined by gold standard FSC=0.143 criterion. Map-to-Model FSC was calculated between the refined atomic models and maps (red line).

**Figure S3.**
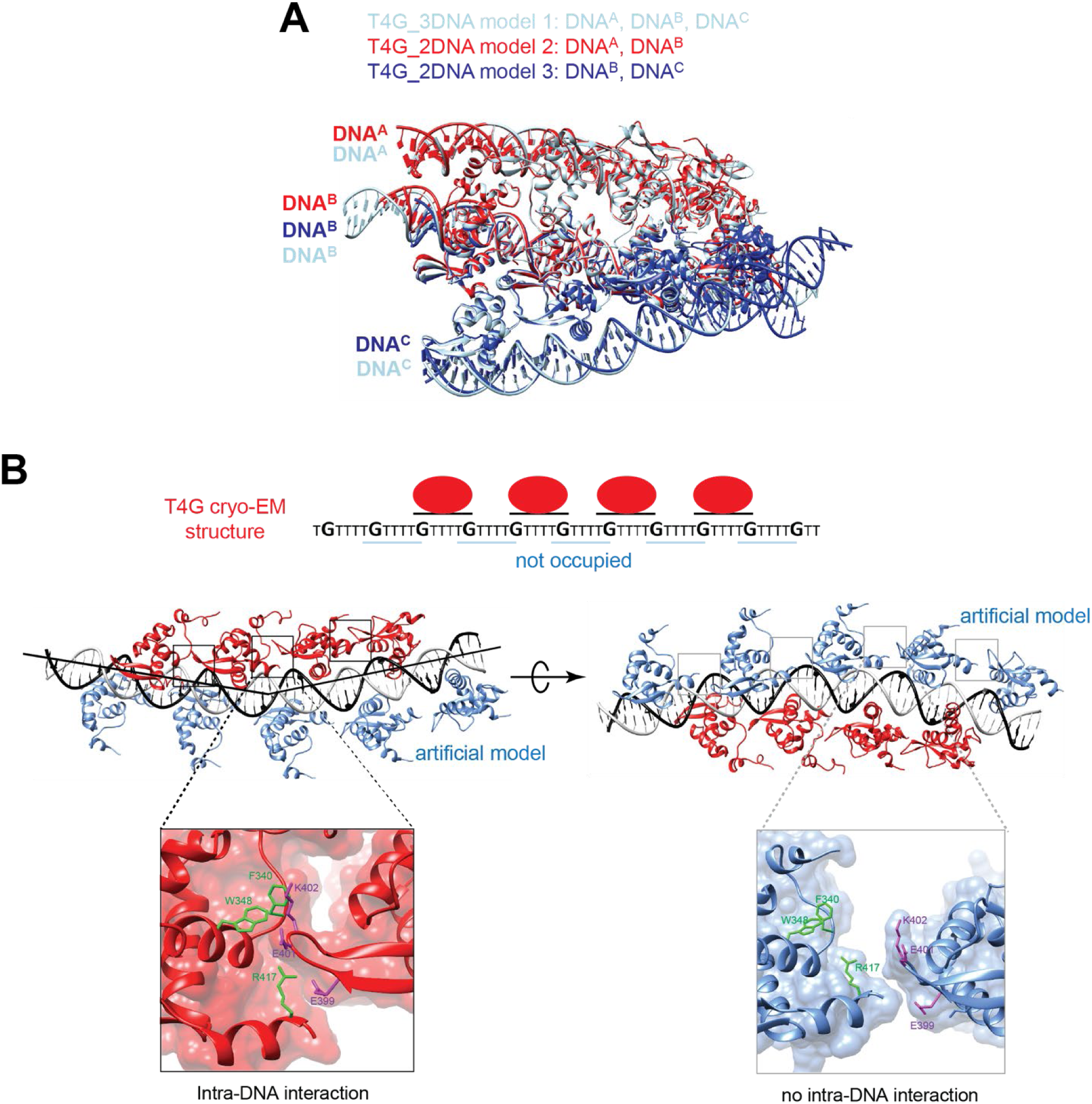
Analysis of FoxP3 multimer structures in complex with T4G repeat DNA. A. Alignment of three different FoxP3^ΔN^–(T_4_G)_15_ complex structures. Model 1 contains three copies of DNA (DNA^A-C^), whereas models 2 and 3 contain two copies of DNA (DNA^A,B^ or DNA^B,C^). Model 1 can be reconstituted by overlaying models 2 and 3 by aligning DNA^B^. B. The structure of FoxP3^ΔN^–(T_4_G)_15_ complex showed that FoxP3 molecules (red) line up along one side of DNA, binding TGTTTTG and forming intra-DNA interactions and bending the DNA towards the bound side. The opposite side remains unoccupied despite having fully accessible binding sites with the same TGTTTTG sequence (blue lines). To understand why FoxP3 occupies only one side of DNA, we modeled in FoxP3 subunits at unoccupied TGTTTTG sites, which revealed a gap between adjacent subunits. This suggests that FoxP3 is able to form intra-DNA interactions only on one side, explaining its occupancy only on one side. Inset: surface representation of adjacent subunits to demonstrate presence and absence of intra-DNA interaction. The interface residues are shown in green and purple.

**Figure S4.**
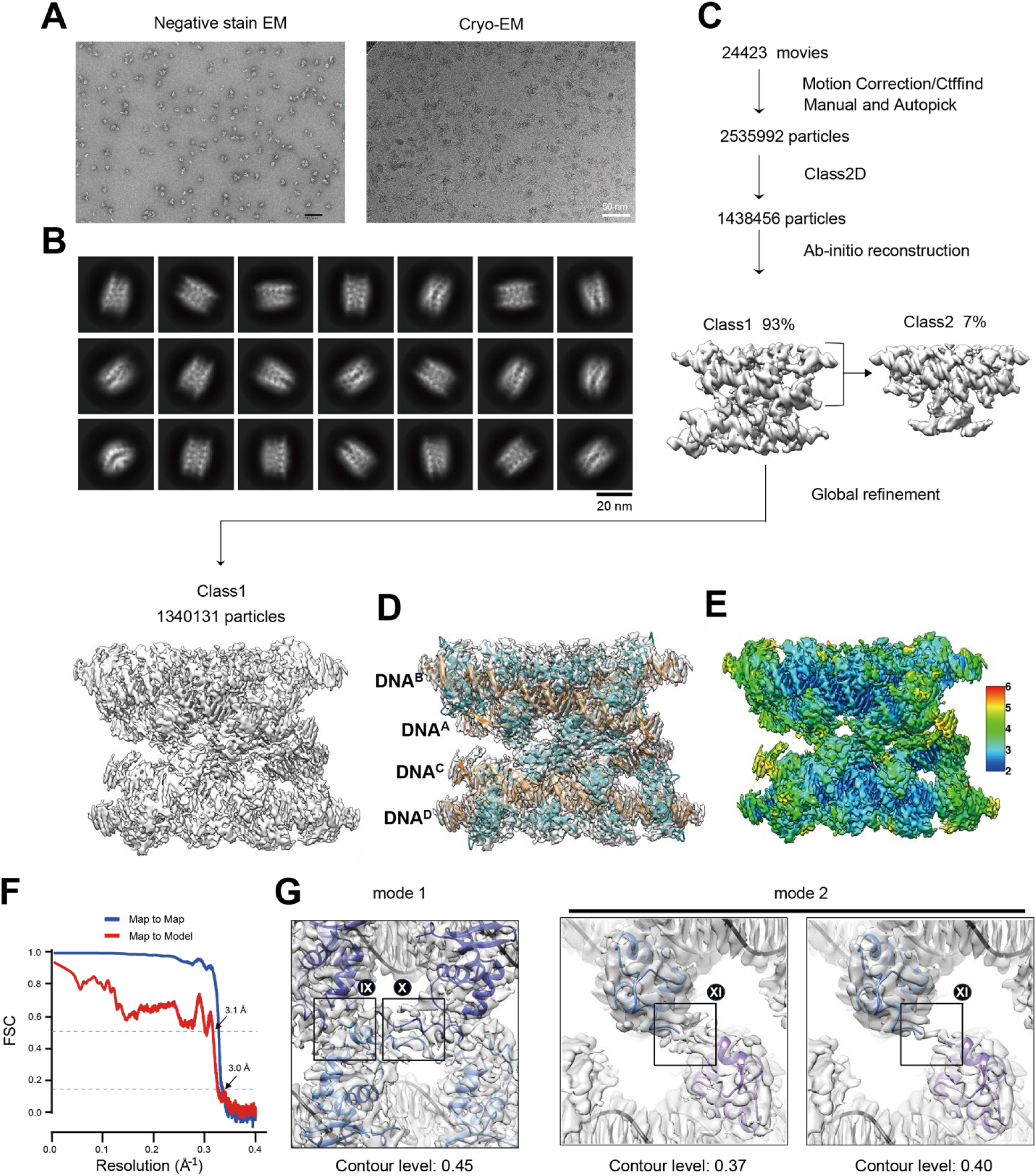
Cryo-EM structure of the FoxP3^ΔN^–(T_2_G)_24_ complex. A. Representative negative-stain EM (left) and cryo-EM images (right) of FoxP3^ΔN^ multimers on (T_2_G)_24_ DNA. B. 2D classes chosen for 3D reconstruction. C. Cryo-EM image processing workflow. See details in Methods. D. Cryo-EM maps and ribbon models of FoxP3^ΔN^ multimer in complex with four copies of (T_2_G)_24_ DNAs. DNA molecules are colored orange. FoxP3^ΔN^ subunits are colored sky blue. E. Local resolution of Cryo-EM map was calculated by CryoSPARC. Resolution range was indicated according to the color bar. F. Fourier shell correlation (FSC) curve. Map-to-Map FSC curve was calculated between the two independently refined half-maps after masking (blue line), and the overall resolution was determined by gold standard FSC=0.143 criterion. Map-to-Model FSC was calculated between the refined atomic models and maps (red line). G. Left: Cryo-EM density map of mode 1 inter-DNA interfaces (IX and X) at 0.45 contour level. Right: Cryo-EM density map of mode 2 inter-DNA interfaces (XI) at 0.37 and 0.40 contour level.

**Figure S5.**
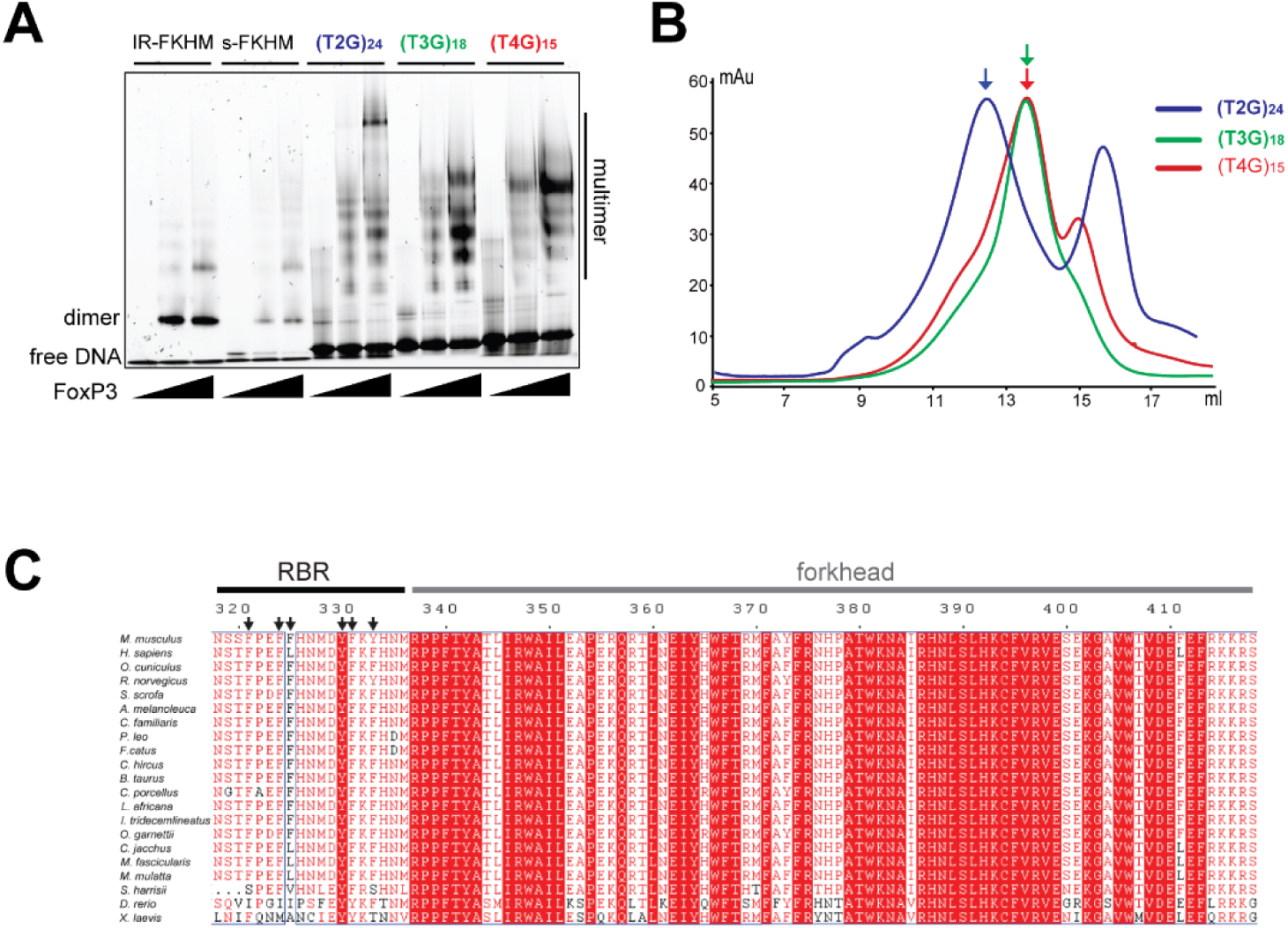
EMSA and SEC analysis of FoxP3-TnG repeat complexes. A. Native gel shift assay of FoxP3^ΔN^ (0, 0.8, 1.6 μM) with DNA harboring (T_2_G)_24_ (72 bp, 0.2 μM), (T_3_G)_18_ (72 bp, 0.2 μM) and (T_4_G)_15_ (75 bp, 0.2 μM). B. Size-exclusion chromatograms (SEC) of FoxP3^ΔN^ in complex with (T_2_G)_24_ (blue), (T_3_G)_18_ (green) and (T_4_G)_15_ (red) after incubated at the 8 (FoxP3^ΔN^) to 1 (DNA) molar ratio. Superpose 6 increase column was used. FoxP3^ΔN^**-**(T_2_G)_24_ complex eluted earlier than FoxP3^ΔN^**-**(T_3_G)_18_ and FoxP3^ΔN^**-**(T_4_G)_15_ complex, suggesting larger complex size. C. Sequence alignment of FoxP3 orthologs from 22 different species. Arrows indicate aromatic residues in the RBR loop of mouse FoxP3. There are 6 aromatic residues within the surface-exposed 17 RBR amino acids, the frequency far greater than the typical frequency of aromatic residues on protein surface.

**Figure S6.**
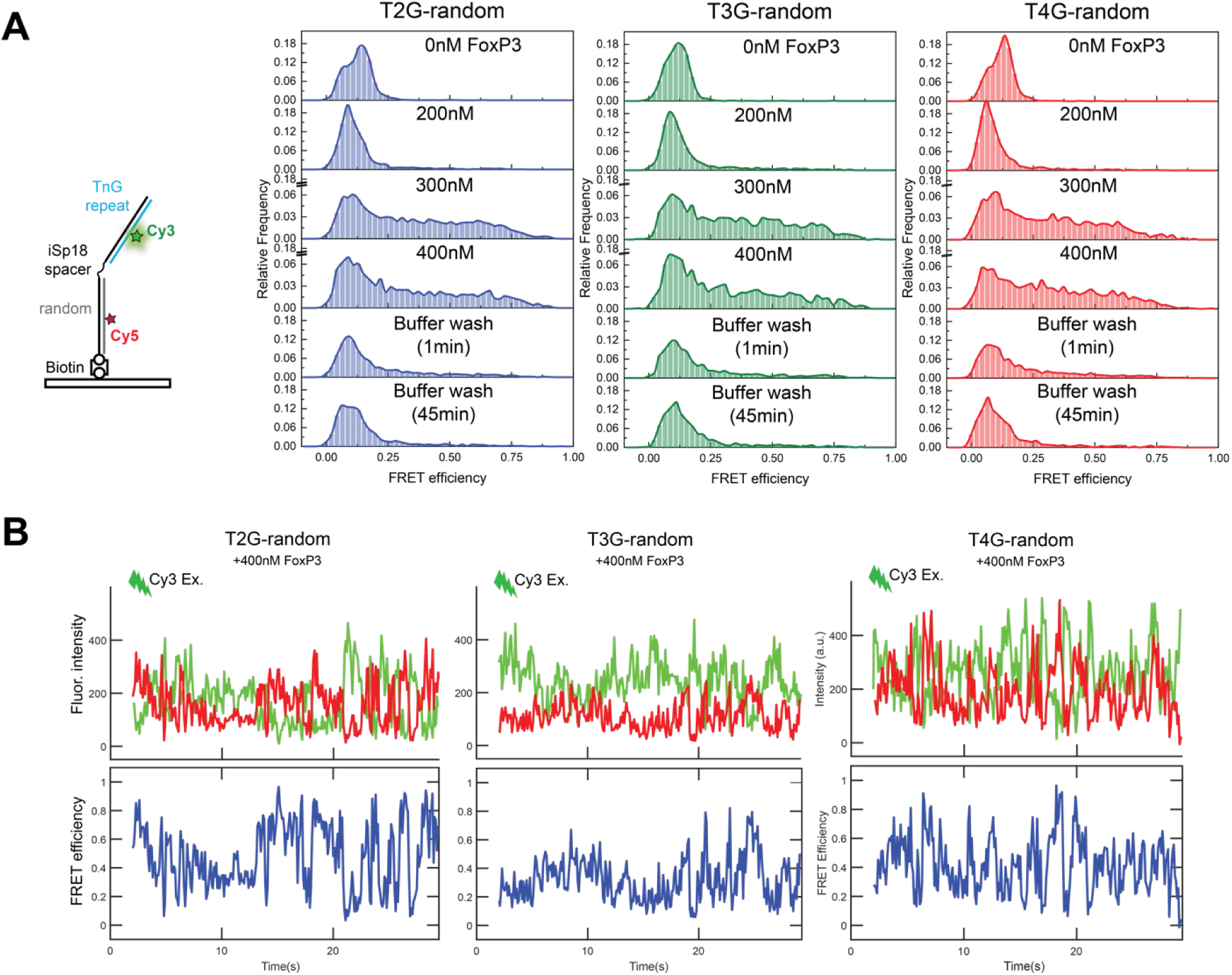
FoxP3-mediated DNA bridging is sequence-dependent. A. Left: TnG-random nunchuck DNA construct used in the experiment. Right: FRET distributions across different TnG-random sequences. Unlike TnG-TnG nunchucks, FRET did not increase at 200 nM FoxP3. While FRET levels rose at higher FoxP3 concentrations (300-400 nM), the increase was lost within 1 min of buffer washing. B. Representative time traces of fluorescent intensity and Cy3-Cy5 FRET efficiency for T2G-random (reproduced in Figure 4D), T3G-random and T4G-random, all in the presence of 400 nM FoxP3.

**Figure S7.**
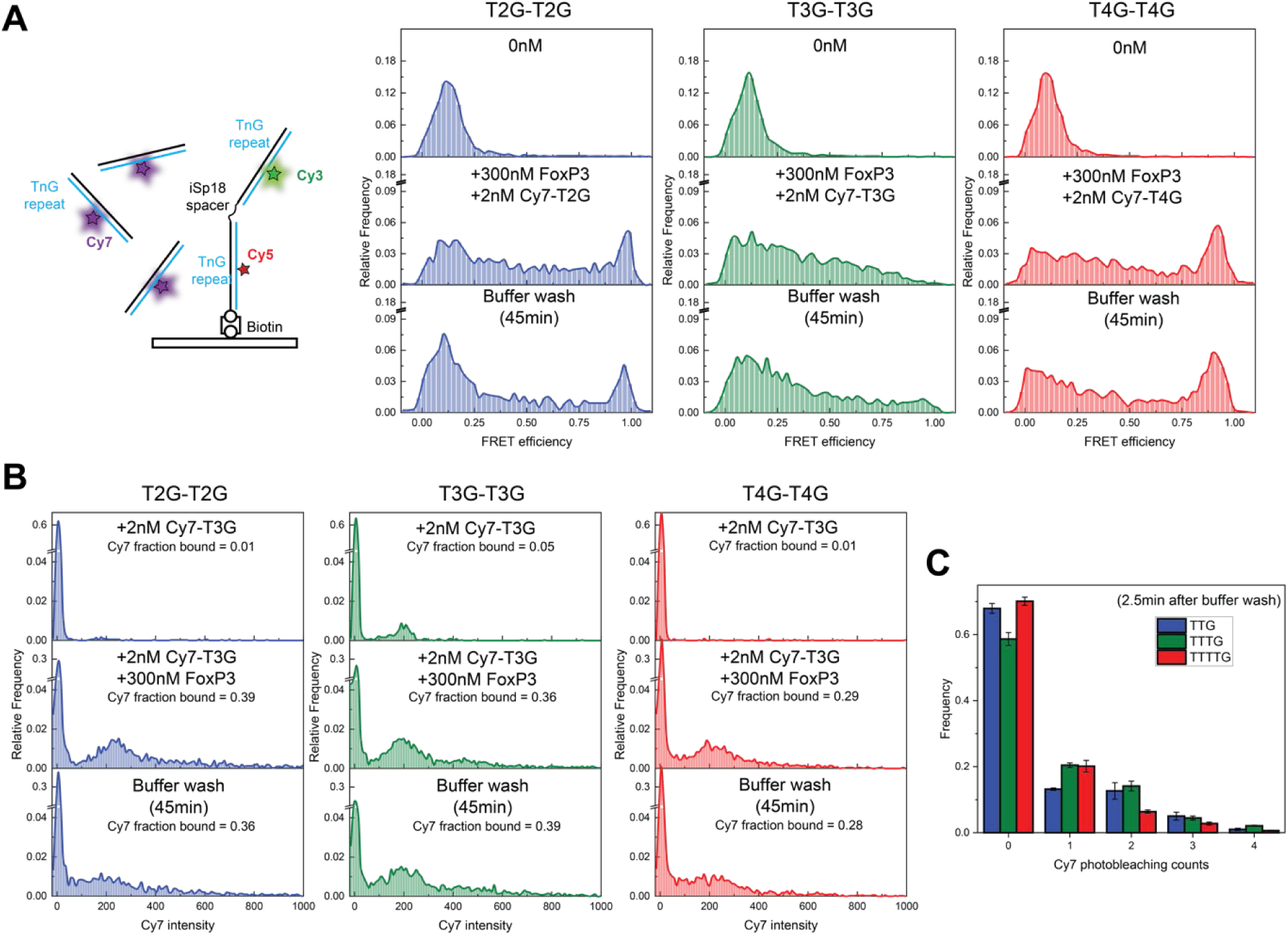
FoxP3-mediated multi-DNA bridging. A-B. Left: experimental setup to examine multi-way DNA bridging by FoxP3 (as in Figure 4E). Cy7-DNA containing TnG repeats was added to immobilized nunchucks along with FoxP3 (300 nM), and were washed with buffer lacking FoxP3 for 45 min. FoxP3-induced FRET (A) and recruitment of Cy7-DNA (B) were measured. C. Cy7 photobleaching counts calculated from single-molecule fluorescent time traces obtained 2.5 min after washing away free FoxP3 and Cy7 DNA.

**Figure S8.**
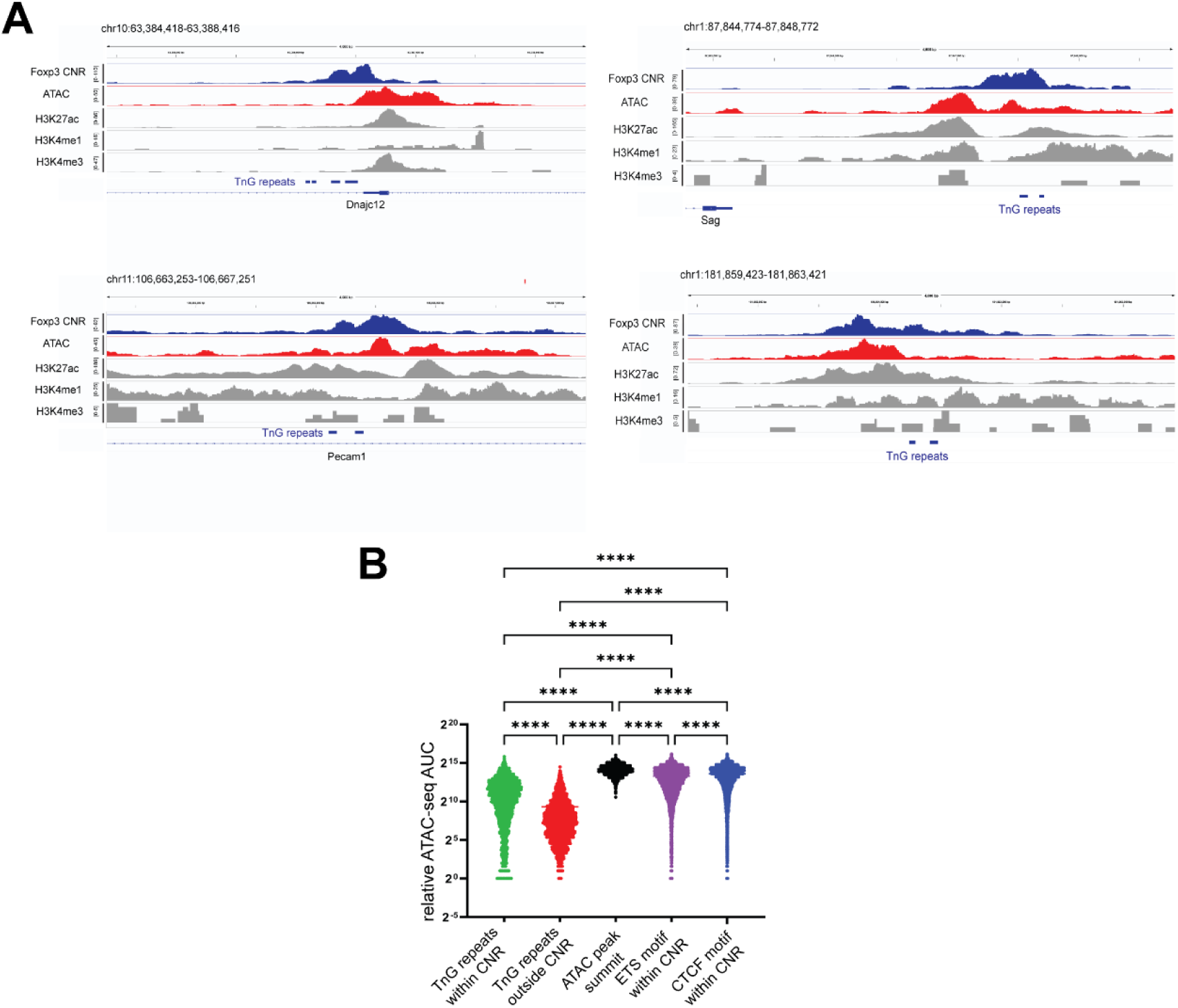
FoxP3-occupied TnG repeats are at the edges of ATAC peaks. A. Representative TnG repeat regions within Foxp3 CNR peaks in genome browser view. Tracks (upper to lower): merged Foxp3 CNR of the Rudensky (PMID:33176163) and Dixon-Zheng datasets (PMID:37932264), thymic Treg ATAC-seq (PMID: 27992401), thymic Treg H3K27ac ChIP-seq (PMID: 27992401), thymic Treg H3K4me1 ChIP-seq (PMID: 27992401), thymic Treg H3K4me3 ChIP-seq (PMID: 27992401), TnG regions within Foxp3 CNR peaks, and Refseq gene annotation. B. Quantitation of (D). ATAC-seq Area under Curve (AuC) within each 600 bp window was calculated and plotted. Statistical analysis was performed using one-way ANOVA; \*\*\*\**P* < 0.0001.

**Figure S9.**
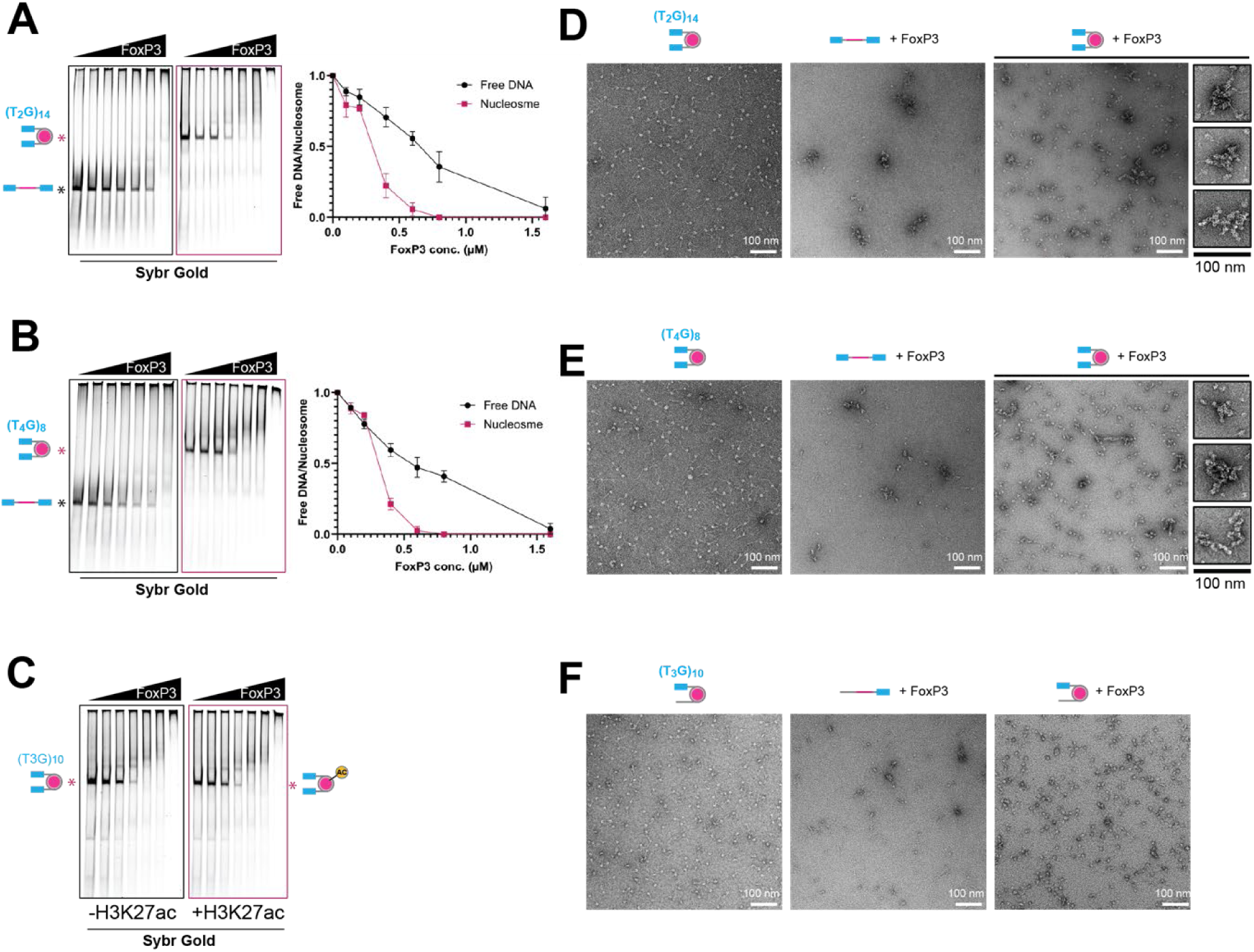
Nucleosomes can facilitate FoxP3’s ability to bridge T_2_G and T_4_G repeats. A. FoxP3 binding to free DNA and nucleosomal DNA harboring (T_2_G)_14_-601-(T_2_G)_14_ sequence as measured by native gel-shift assay. Free DNA and nucleosome (50 nM each) were incubated with an increasing concentration of FoxP3 (0, 0.1,0.2,0.4, 0.6, 0.8, 1.6 μM). Sybrgold stain was used for visualization and quantitation for FoxP3-free species of DNA. B. Same as (A), but using DNA harboring (T_4_G)_8_-601-(T_4_G)_8_. C. Comparison of FoxP3 binding to nucleosomal DNA harboring (T_3_G)_10_-601-(T_3_G)_10_, with and without H3K27ac. D-F. Representative negative stain electron micrographs of nucleosomal DNA without FoxP3 (left), nucleosome-free DNA with FoxP3 (middle), and nucleosomal DNA with FoxP3 (right). DNA has the (T_2_G)_14_-601-(T_2_G)_14_ sequence in (D), (T_4_G)_8_-601-(T_4_G)_8_ in (E) and (T_3_G)_10_-601-random in (F). The random sequence in (F) is 40 bp. Aggregated particles (black boxes) were highly enriched in FoxP3-bound nucleosomal DNA in (D) and (E), but not in (F).

### KEY RESOURCES TABLE

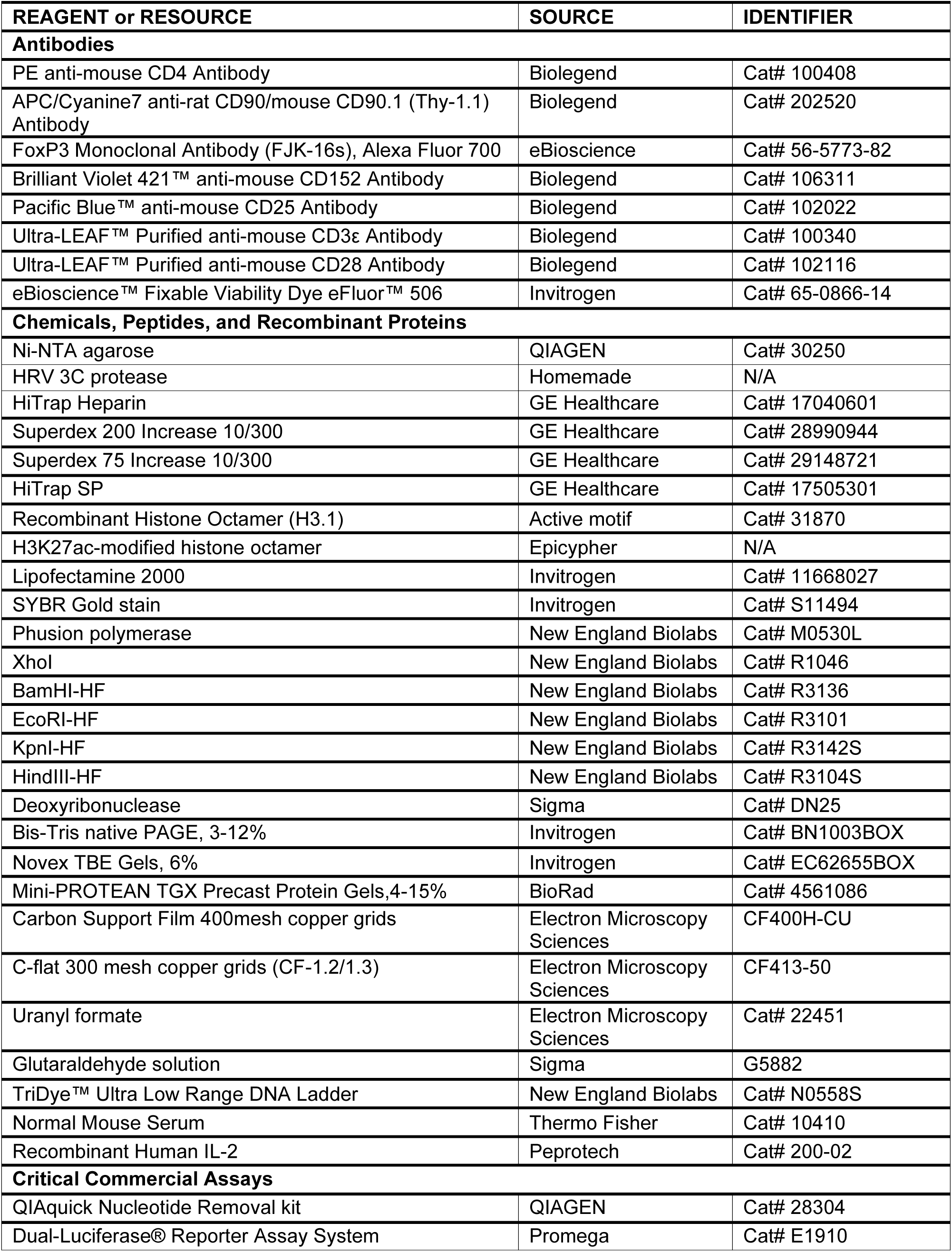
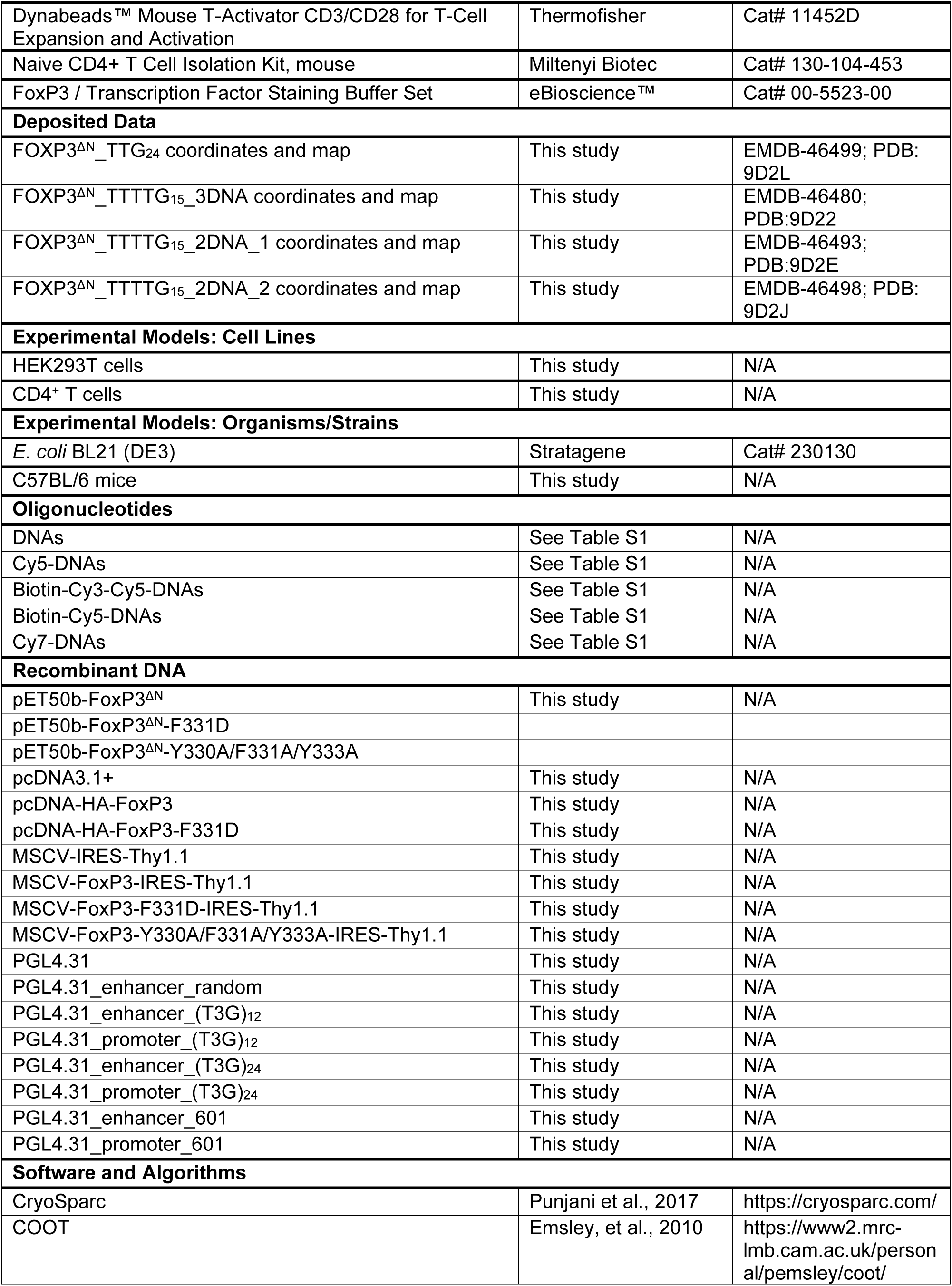
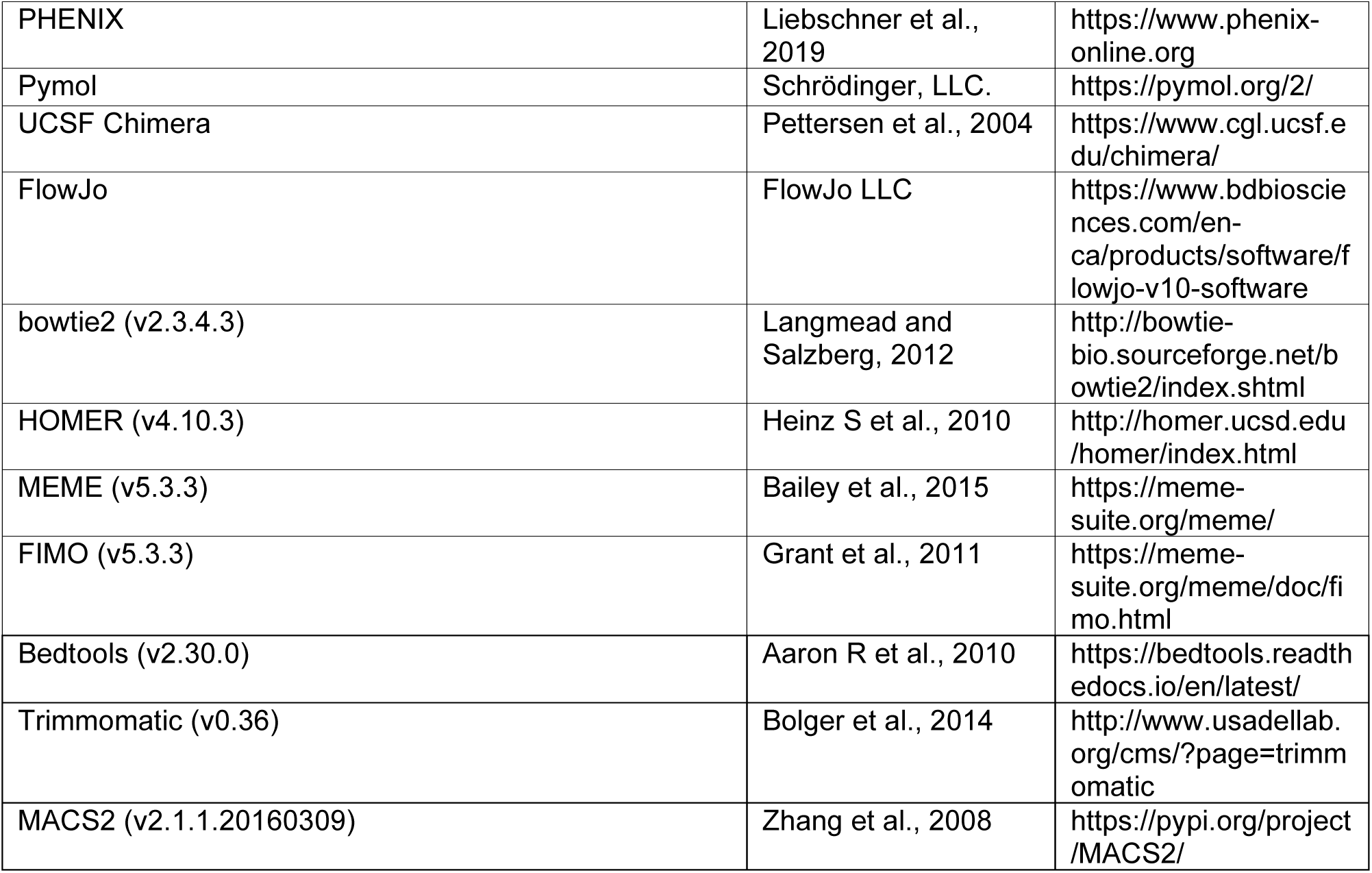

### RESOURCE AVAILABILITY

#### Lead contact

Further information and requests for resources and reagents should be directed to and will be fulfilled by the Lead Contact, Sun Hur

#### Materials Availability

All plasmids generated in this study are available from the Lead Contact with a completed Materials Transfer Agreement.

#### Data and Code Availability

The CryoEM volumes have been deposited in the Electron Microscopy Data Bank (EMDB) with accession codes: 46499, 46480, 46493, 46498. The atomic coordinates have been deposited in the Protein Data Bank with accession codes: 9D2L, 9D22, 9D2E, 9D2J.

### EXPERIMENTAL MODEL AND SUBJECT DETAILS

#### Mice

C57BL/6N mice, sourced from Taconic Biosciences, were housed in an individually ventilated cage system at the specific-pathogen-free New Research Building facility of Harvard Medical School. The mice were maintained at a controlled environment with a temperature of 20-22°C, humidity ranging from 40-55%, and a 12-hour light-dark cycle. The spleens of 12∼14 weeks old female C57BL/6 mice were isolated for the study.

#### Naive CD4^+^ T Cells

Cells were isolated by using Naive CD4^+^ T Cell Isolation Kit (Miltenyi Biotec, Cat#130-104-453) according to the manufacturer’s instructions and maintained in complete RPMI medium (10% FBS heat-inactivated, 2mM L-Glutamine, 1mM Sodium Pyruvate, 100μM NEAA, 5mM HEPES, 0.05mM 2-ME).

#### HEK293T

Cells were maintained in DMEM (High glucose, L-glutamine, Pyruvate) with 10% fetal bovine serum.

### METHOD DETAILS

#### Material Preparation Plasmids

Mouse FoxP3 plasmids were made as previously described^1^. For Mammalian expression plasmids, HA-tagged mouse FoxP3 CDS was inserted into pcDNA3.1+ vector between KpnI and BamHI sites. For bacterial expression plasmids, the genes encoding mouse FoxP3^ΔN^ (residues 188-423) was inserted into pET50b between Xmal and HindIII sites. For retroviral packaging plasmids, HA-tagged mouse FOXP3 CDS was inserted into MSCV-IRES-Thy1.1 vector. For luciferase assay plasmids, different combinations of T3G repeat sequence, 601 sequence, Gal4 UAS and Ad promoter were ordered as gblocks, and inserted into PGL4.31 plasmids between KpnI and HindIII. For enhancer plasmids used in luciferase assay, stop code was introduced at the 6^th^ amino acid (N) of luciferase to interrupt the expression. All mutations within FoxP3 were generated by site-directed mutagenesis using Phusion High Fidelity (New England Biolabs) DNA polymerases.

#### DNA oligos

Single-stranded DNA (ssDNA) oligos were synthesized by IDTDNA. Double-stranded DNA (dsDNA) oligos for nucleosome reconstitution, negative staining EM and EMSA assay were annealed from single-stranded, complementary oligos. After briefly spinning down each oligonucleotide pellet, ssDNAs were dissolved in the annealing buffer (10 mM Tris-HCl pH 7.5, 50 mM NaCl). Complementary ssDNAs were then mixed in equal molar amounts, heated to 94°C for 2 minutes and gradually cooled down to room temperature. For dsDNA in SEC and cryo-EM analysis, HPLC purified single-stranded, complementary oligos were purchased from IDTDNA. After annealing, dsDNA was further purified by size-exclusion chromatography on Superdex 75 Increase 10/300 (GE Healthcare) columns in 20 mM Tris-HCl pH 7.5, 150 mM NaCl. Cy5 labeled ssDNA oligo was synthesized by IDTDNA and dissolved in annealing buffer (10 mM Tris-HCl pH 7.5, 50 mM NaCl), and then mixed together with complementary unlabeled ssDNA in equal molar amounts, heated to 94°C for 2 minutes and gradually cooled down to room temperature. DNA oligonucleotides for single-molecule FRET were purchased from Integrated DNA Technologies (IDT). Amine-modified oligos were labeled with either Cy3, Cy5, LD655 or Cy7 by mixing 80µM oligo with 4mM NHS-dye, and 200mM fresh NaHCO3 in a 62.5µl reaction volume for 4h at room temperature, followed by overnight incubation at 4°C. Free dye was removed using ethanol precipitation. DNA oligos for TnG-TnG and TnG-random nunchuck DNA constructs were annealed by mixing the three strands (TnG-TnG backbone, TnG 1 and TnG/random Biotin) with a molar ratio of 1:1.6:0.5 in 20µl of 10mM Tris:HCl (pH 8.0) and 50mM NaCl (T50 buffer). The annealing reaction was performed by slow cooling from 95°C to 25°C for over an hour in a thermocycler. TnG LD655 and TnG Cy7 was prepared by mixing the two strands (TnG/random backbone and TnG 1/random 1) with the molar ratio of 1:1.2, followed by annealing using the slow cooling cycle. Annealing efficiency for all constructs was assayed using a 6% native PAGE. Sequence of all the DNA oligos used are shown in Extended Data Table 1.

#### TnG repeat sites within or outside FoxP3 CNR peaks

The union of Foxp3 CNR peaks were acquired by combining the Treg Foxp3 CNR peaks from Sasha group^2^ (GSE154680) and Dixon group^3^ (GSE217147). Treg ATAC peaks outside Foxp3 CNR peaks were acquired by combining Treg ATAC-seq peaks from ImmGen, and then removing all regions within 10kb distance of any combined CNR peaks. TnG motifs were used as FIMO^4^ search inputs for finding TnG regions within Foxp3 CNR peaks or Treg ATAC peaks outside Foxp3 CNR peaks. FIMO search outputs were filtered by [p-value < 8e-5], and [no consecutive 6TGs, 6TAs, 6TCs, 6GAs, 6GCs, 6Acs, 12Ts, 12Gs, 12Cs or 12As]. The filtered FIMO regions were merged into TnG repeat sites within (n=2,019) or outside FoxP3 CNR peaks (n=3,579).

#### TnG 1kb clustering quantification

To quantify the clustering of different TnG groups, the center coordinates of TnG regions were calculated, and expanded up to +/-500bp to make a region of 1kb total. Of these 1kb regions, TnG regions were searched by FIMO^4^ within Foxp3 CNR peaks and Treg ATAC peaks outside of Foxp3 CNR peaks, and filtered using the same criteria described in the method above. The total number of TnG bps within 1kb was then calculated.

#### Foxp3 RNA-seq quantification

RNA-seq raw datasets of Treg and Tcon were downloaded from GSE154680^2^. The fastq files were paired, trimmed with Trimmomatic^5^ (v0.36), and mapped to mm10 reference genome using STAR aligner^6^ (v2.7.0a). Exon reads were counted by R library Count_feature from Rsubread^7^ (v4.4), and multi-mapping reads were counted by fraction.

#### Protein expression and purification

All recombinant proteins in this paper were expressed in BL21(DE3) at 18°C for 16-20 hrs following induction with 0.2 mM IPTG. Cells were lysed by high-pressure homogenization using an Emulsiflex C3 (Avestin). All proteins are from the *Mus. musculus* sequence, unless mentioned otherwise. FoxP3^ΔN^ (residues 188-423) was expressed as a fusion protein with an N-terminal His_6_-NusA tag. After purification using Ni-NTA agarose, the protein was treated with HRV3C protease to cleave the His_6_-NusA-tag and were further purified by a series of chromatography purification using HiTrap Heparin (GE Healthcare), Hitrip SP (GE Healthcare) and Superdex 200 Increase 10/300 (GE Healthcare) columns. The final size-exclusion chromatography (SEC) was done in 20 mM HEPES pH 7.5, 500 mM NaCl, 2 mM DTT.

#### Cryo-EM sample preparation and data collection

FoxP3^ΔN^ was incubated with (T_4_G)_15_ and (T_2_G)_24_ DNA at a molar ratio of 8:1 in buffer 20 mM HEPES pH 7.5, 150 mM NaCl, 2 mM DTT at RT for 10 mins. The complex was then crosslinked using 0.05 % glutaraldehyde for 10 mins at RT prior to quenching with 1/10 volume of 1M Tris-HCl pH 7.5 (for a final Tris concentration of 0.1 M). The FoxP3^ΔN^-DNA complex was then purified by Superose 6 Increase 10/300 GL (GE Healthcare) column in 20mM Tris-HCl pH 7.5, 100 mM NaCl, 2 mM DTT. The sample was concentrated to 1 mg/ml (final for protein) and applied to freshly glow-discharged C-flat 300 mesh copper grids (CF-1.2/1.3, Electron Microscopy Sciences) at 4℃. The grids were plunged into liquid ethane after blotting for 5 s using Vitrobot Mark IV (FEI) at the humidity setting of 100%. The grids were screened at the Harvard Cryo-EM Center and UMass Cryo-EM core facility using Talos Arctica microscope (FEI). The grids that showed a good sample distribution and ice thickness were used for data collection on Titan Krios (Janelia Cryo-EM facility) operated at 300 kV and equipped with Gatan K3 camera. 14535 micrographs of FoxP3^ΔN^–(T_4_G)_15_ complex and 24423 micrographs of FoxP3^ΔN^–(T_2_G)_24_ complex were taken at a magnification of 105,000x with a pixel size of 0.827 Å. Each movie comprised of 60 frames at total dose of 60 e^-^/Å^2^. The data were collected in a desired defocus range of -0.7 to -2.1 μm.

#### Cryo-EM data processing and structure refinement

Data were processed using cryoSPARC V4.2.0^8^. Movies were motion corrected in CryoSparc followed by patch CTF estimation. A set of templates were then generated by blob picker and used for template-based autopick in CryoSparc. For FoxP3^ΔN^–(T_4_G)_15_ complex, 1327974 raw particles were used for 2D classification. 1086082 particles from selected 2D classes were used for Ab-initio reconstruction, where they were divided into 3 Ab-initio classes. 630524 particles from class 1, 195660 particles from class 2 and 259858 particles from class 3 were refined to a final resolution of 2.6 Å (map I), 2.9 Å (map II), 2.8 Å (map III) respectively with non-uniform refinement. For structure refinement, a previous crystal structure of a FoxP3^ΔN^ monomer bound to DNA (PDB: 7TDX) was docked into the EM density map using UCSF Chimera^9^. A total of 11 copies of FoxP3^ΔN^ monomers with DNA were docked into map I, 8 copies of FoxP3^ΔN^ monomers with DNA were docked into map II and map III. Subsequently, three different models were built manually against the respective EM density map using COOT^10^, and refined using phenix.real_space_refine^11^. The structure validation was performed using MolProbity^12^ from the PHENIX package. The curve representing model versus full map was calculated, based on the final model and the full map. All molecular graphics figures were prepared with UCSF Chimera^9^. All softwares used for cryo-EM data processing and model building were installed and managed by SBGrid^13^. For FoxP3^ΔN^–(T_2_G)_24_ complex, 2535992 raw particles were used for 2D classification. 1438456 particles from selected 2D classes were used for Ab-initio reconstruction, where we got two Ab-initio classes, class 1 with 1340131 particles and class 2 with 98325 particles. Class 1 particles were further refined to a final resolution of 2.6 Å with non-uniform refinement. A total of 28 copies of FoxP3^ΔN^ monomers with DNA were docked into the density map, the following model building and refinement is the same as the method described above. The statistics of the 3D reconstruction and model refinement are summarized in Extended Data Table 2.

#### Electrophoretic Mobility Shift Assay (EMSA)

TnG repeats DNA (0.2μM) was mixed with the indicated amount of FoxP3^ΔN^ and mutations in the EMSA buffer A (20mM HEPES pH 7.5, 150mM NaCl, 1.5mM MgCl_2_ and 2mM DTT), incubated for 30 min at 4 °C and analyzed on 3-12% gradient Bis-Tris native gels (Life Technologies) at 4 °C. After staining with Sybr Gold stain (Life Technologies), Sybr Gold fluorescence was recorded using ChemiDoc Imaging Systems (Bio-Rad) and analyzed with ChemiDoc Analysis Software.

#### CD4_+_ T Cell isolation and retroviral transduction

Naïve CD4^+^ T cells were isolated by negative selection from mouse spleens using the isolation kit (Miltenyi Biotec) according to the manufacturer’s instruction. The purity was estimated to be >90% as measured by PE anti-CD4 (Biolegend) staining and FACS analysis. Naïve CD4^+^ T cells were then activated with anti-CD3 (Biolegend), anti-CD28 (Biolegend) and 50 U/mL of IL2 (Peprotech) in complete RPMI medium (10% FBS heat-inactivated, 2 mM L-Glutamine, 1 mM Sodium Pyruvate, 100 μM NEAA, 5 mM HEPES, 0.05 mM 2-ME). The activation state of T cells was confirmed with increased cell size and CD44 (BioLegend) expression by FACS. After 48 hours, cells were spin-infected with retrovirus containing supernatant from HEK293T cells transfected with retroviral expression plasmids (Empty MSCV-IRES-Thy1.1 vector, wildtype-FoxP3 and mutations encoding vectors) and cultured for 2∼3 days in complete RPMI medium with 100 U/mL of IL2.

#### FoxP3 transcriptional activity assay in CD4_+_T cells

FoxP3 transcriptional activity was measured by levels of known target gene CTLA4, and the FoxP3 expression marker Thy1.1. FoxP3-transduced CD4^+^ T cells were stained with antibody targeting Thy1.1 (Biolegend) on day 2 post-retroviral infection. The level of CTLA4 was measure by intracellular staining using anti-CTLA4 (Biolegend) and the Transcription Factor Staining Buffer Set (eBioscience) on day 3 post retroviral infection. Flow cytometry data were analyzed with FlowJo software and presented as plots of mean fluorescence intensity (MFI) of CTLA4 in cells grouped into bins of Thy1.1 intensity, which is the expression marker for FoxP3. Each result is representative of 3 independent experiments.

#### Single molecule imaging setup and preparation

smFRET experiments were performed using a prism-based total internal reflection fluorescence (TIRF) microscope. Fluorophores were excited by solid-state lasers: 640nm - Coherent (Cy5 and LD655), 543nm (Cy3) and 750nm (Cy7) - Shanghai Dream Lasers Technology. Fluorescence signal was collected by a Nikon water immersion 60x/1.27 NA objective and a custom laser-blocking filter. Emissions were separated into two or three channels using dichroic mirrors. Images were captured using an electron-multiplying charge-coupled device (Andor iXon 897). Parameters for spot detection, background correction, donor leakage and Cy3-Cy5 crosstalk correction were calculated as discussed previously^14^ using custom IDL (Interactive Data Language) and MATLAB scripts (https://sites.google.com/site/taekjiphalab/resources). The methoxy polyethylene glycol (PEG) passivated quartz slides containing biotin-PEG and coverslips were purchased from Nano Surface Sciences and assembled into flow channels. Neutravidin (0.2 mg/ml in T50 buffer) was flown into each channel to prepare a functionalized surface by binding to biotinylated PEG from the quartz slide. After 1 min of incubation, the neutravidin solution was washed off using T50 buffer and the channel was used for DNA immobilization.

#### Two-color DNA nunchuck smFRET imaging

All fluorescently labeled pre-annealed TnG-TnG or TnG-random DNA constructs were diluted to 50pM in FoxP3 buffer composed of 20mM Tris (pH7.5), 150mM NaCl, 2mM MgCl2, 0.1mg/ml BSA. DNA was incubated for 2 min on the neutravidin-functionalized chamber. Free molecules were washed out with FoxP3 buffer. Imaging buffer composed of FoxP3 buffer along with an oxygen-scavenging system [0.8% (w/v) dextrose, 2mM Trolox, glucose oxidase (1mg/ml, Sigma-Aldrich), and catalase (500U/ml, Sigma-Aldrich)] was flown to the channel. A total of 10 short movies (100ms exposure time) were taken to capture the fluorescent signal of surface immobilized DNA prior to the addition of FoxP3 protein. 20 frames were recorded per movie, 10 frames were taken using Cy3 excitation, followed by 10 frames of Cy5 excitation. FRET histograms were built using similar imaging settings before and after protein addition. Only molecules having both Cy3 and Cy5 were selected, and two-color FRET efficiencies were calculated as previously described^14^. FoxP3 and mutant FoxP3 F331D were diluted to the desired concentrations using imaging buffer and flown into the chamber. Each protein condition was incubated for 10 min before imaging. Short movies were taken as explained above. Long movies were recorded for a total of 1000 frames using a 50ms exposure time. FRET efficiency time trajectories were calculated from fluorescent intensities of single molecules. Free FoxP3 protein was washed out using imaging buffer and short movies were taken over time to measure the stability of FoxP3 bridging DNA.

#### Three-color colocalization single-molecule assay

Cy3-Cy5 fluorescently labeled TnG-TnG DNA constructs were immobilized as described above. To measure the ability of FoxP3 bridging more than two copies of TnG DNA, 2nM of Cy7 labeled non-biotinylated TnG DNA and FoxP3 (300nM) were added simultaneously. Free Cy7- TnG or Cy7-random and FoxP3 were washed off after 10 min of incubation, and a total of 10 short movies were taken using alternating laser excitation (ALEX). Cy3, Cy5 and Cy7 were individually excited for 200ms each over a total of 30 frames. Spots that had both Cy3 and Cy5 signal were selected and the Cy3-Cy5 FRET efficiency was calculated together with the fraction of Cy7 molecules that colocalized (bound) to these dual labeled immobilized DNA. Long movies with a total of 1500 frames (10 frames-Cy3 excitation, 10 frames-Cy5 and the remaining frames being Cy7 excitation) were taken to photobleach and quantify the Cy7-TnG molecules bound to the nunchuck DNA. To account for unspecific colocalization of Cy7-DNA to Cy3-Cy5 immobilized nunchuck DNA, we measured the fraction of Cy7 molecules in the absence of FoxP3.

#### Pull-down experiments for stoichiometry analysis

To quantify the DNA multimers bridged by FoxP3, we incubated biotin LD655 pre-annealed TnG DNA (5nM) with 1µM of FoxP3 along with FoxP3 buffer in an Eppendorf tube (Protein Lobind) for 10 min. Next, the reaction was diluted 150 times and flown into the neutravidin-functionalized chamber. After 10 min of incubation, unbound DNA-FoxP3 multimers were washed off with FoxP3 buffer. Imaging buffer without glucose oxidase (for complete bleaching of labeled molecules) was added. Long movies were recorded by directly exciting LD655 for 1000 frames using an exposure time of 100ms. The increased photostability of LD655 fluorophore is used to precisely quantify the photobleaching steps across the different TnG constructs.

#### ATAC_tTreg AuC intensity and heatmap

The summits of ATAC thymus Treg^15^ (SRR5385307) were first computed using MACS2^16^ (v2.1.1.20160309), and then filtered for only the regions within Foxp3 CNR peaks. The CTCF and ETS1 motifs were downloaded from MEME JASPAR 2022 database^17^, and searched by FIMO within Foxp3 CNR peaks with a p-value cutoff of 0.01. The overlapping FIMO regions were merged with bedtools merge. Then, the centers of TnG within Foxp3 CNR, ATAC tTreg summits, CTCF and ETS1 motif centers were all expanded +/-300bp, and heatmaps were drawn with ATAC_tTreg intensity by deeptools^18^ (v3.0.2). For each individual heatmap, the regions were sorted by ATAC_tTreg intensity in descending order.

#### Distance from ATAC summit to closest motif center

The center coordinates of TnG regions, ETS motifs and CTCF motifs within Foxp3 CNR peaks were calculated by the mean value of start and end coordinates. Then, the distances from ATAC summits to the closest center of TnG region, ETS motif or CTCF motif were calculated by bedtools closest (v2.30.0) for up to 5kb.

#### Nucleosome reconstitution and EMSA analysis

Nucleosome core particles (NCP) were reconstituted with recombinant Histone Octamer H3.1 (Active motif) and DNAs harboring 601 sequence as described previously^19^. Briefly, 1 μM of DNAs were incubated with 1 μM of the histone octamer and were dialyzed against 10mM Tris-HCl PH 7.5, 1mM EDTA, 2mM DTT for 24 hrs. Nucleosomes (0.05 μM) were incubated with the indicated amount of FoxP3^ΔN^ in the EMSA buffer B (10mM Tris-HCl pH 7.5, 50mM NaCl, 1mM EDTA, and 2mM DTT) for 30 min at 4 °C and analyzed on 6% TBE gels (Life Technologies) at 4 °C. After staining with Sybr Gold stain (Life Technologies), Sybr Gold fluorescence was recorded using ChemiDoc Imaging Systems (Bio-Rad) and analyzed with ChemiDoc Analysis Software.

#### Negative stain EM

Nucleosomes (0.05 μM) were incubated with 0.8 μM FoxP3^ΔN^ in the EMSA buffer B (10mM Tris-HCl pH 7.5, 50mM NaCl, 1mM EDTA, and 2mM DTT) for 30 min at 4 °C. The samples were then diluted 10-fold with the same buffer, immediately adsorbed to freshly glow-discharged carbon-coated grids (Ted Pella) and stained with 0.75% uranyl formate as described before^20^. Images were collected using a JEM-1400 transmission electron microscope (JEOL) at 50,000X magnification.

#### Luciferase assay

Two types of plasmids were generated for luciferase assay: an enhancer plasmid containing an enhancer element (UAS) which can be bound by Gal4 DBD fused with an activation domain from an unrelated TF Aire (Gal4-TAD), and reporter plasmid containing the firefly (FF) luciferase gene driven by a minimal promoter. Both plasmids contained (T_3_G)n repeats (n=12 or 24) or (T_3_G)12-601-(T_3_G)12 sequence, upstream of the promoter or enhancer. 50ng enhancer plasmid and 50ng reporter plasmid, along with 200ng pcDNA plasmid expressing FoxP3 and 10ng Renilla luciferase-encoding transfection control plasmid, were co-transfected into 293T cells (∼80% confluence) stably expressing Gal4-TAD under the control of doxycycline (Dox). 500ng/ml Dox was added at 6 h post-transfection. 18 h later, Gal4DBD-CTT transcriptional activity was measured by Dual Luciferase Reporter assay (Promega) using a Synergy2 plate reader (BioTek). Firefly luciferase activity was normalized against Renilla luciferase activity. Data are representative of three biological replicates.

#### Code availability

The custom codes used in this manuscript can be found in this Github link: https://github.com/DylannnWX/Hurlab_Nucleosome_Foxp3_Manuscript

## Notes

### Competing Interest Statement

The authors have declared no competing interest.

